# TRPC3 suppression ameliorates synaptic dysfunctions and memory deficits in Alzheimer’s disease

**DOI:** 10.1101/2024.09.16.611061

**Authors:** Jiaxing Wang, Ling Chen, Zhengjun Wang, Sicheng Zhang, Dongyi Ding, Geng Lin, Hua Zhang, Vijay K. Boda, Dehui Kong, Tyler C. Ortyl, Xusheng Wang, Lu Lu, Fu-Ming Zhou, Ilya Bezprozvanny, Jianyang Du, Zhongzhi Wu, Wei Li, Francesca-Fang Liao

## Abstract

Transient receptor potential canonical (TRPC) channels are widely expressed in the brain; however, their precise roles in neurodegeneration, such as Alzheimer’s disease (AD) remain elusive. Bioinformatic analysis of the published single-cell RNA-seq data collected from AD patient cohorts indicates that the *Trpc3* gene is uniquely upregulated in excitatory neurons. TRPC3 expression is also upregulated in post-mortem AD brains, and in both acute and chronic mouse models of AD. Functional screening of TRPC3 antagonists resulted in a lead inhibitor JW-65, which completely rescued Aβ-induced neurotoxicity, impaired synaptic plasticity (e.g., LTP), and learning memory in acute and chronic experimental AD models. In cultured rat hippocampal neurons, we found that treatment with soluble β-amyloid oligomers (AβOs) induces rapid and sustained upregulation of the TRPC3 expression selectively in excitatory neurons. This aberrantly upregulated TRPC3 contributes to AβOs-induced Ca^2+^ overload through the calcium entry and store-release mechanisms. The neuroprotective action of JW-65 is primarily mediated via restoring AβOs-impaired Ca^2+^/calmodulin-mediated signaling pathways, including calmodulin kinases CaMKII/IV and calcineurin (CaN). The synaptic protective mechanism via TRPC3 inhibition was further supported by hippocampal RNA-seq data from the symptomatic 5xFAD mice after chronic treatment with JW-65. Overall, these findings not only validate TRPC3 as a novel therapeutic target for treating synaptic dysfunction of AD but most importantly, disclose a distinct role of upregulated TRPC3 in AD pathogenesis in mediating Ca^2+^ dyshomeostasis.

## Introduction

Alzheimer’s disease (AD) is a progressive neurological condition featured pathologically by the extracellular amyloid-β (Aβ)-containing plaques and intracellular tau-laden neurofibrillary tangles, as well as neuronal loss and neuroinflammation. AD manifests significant cognitive decline, which primarily affects memory. Genetic mutations in genes encoding for amyloid precursor protein or presenilin are associated with early onset of AD in humans, resulting in increased Aβ production and disrupted intracellular Ca^2+^ homeostasis. This was the initial fundamental evidence supporting the two dominating hypotheses of AD pathogenesis: the amyloid cascade hypothesis[1, 2] and the Ca^2+^ hypothesis[3, 4]. However, the channels involved in disrupted Ca^2+^ homeostasis are not fully identified. One out of the two FDA-approved AD drugs memantine as a non-competitive antagonist to NMDA receptor/NR2B can only relieve symptoms in mild AD patients, implying that other types of Ca^2+^ channels may also play a significant role in AD pathogenesis.

Ca^2+^ homeostasis is crucial for neuronal functions, which include development, survival, synaptic transmission, memory, and plasticity. Cellular Ca^2+^ signals are regulated by the entry of Ca^2+^ from the extracellular environment or the release of Ca^2+^ from intracellular compartments through receptor/store-operated Ca^2+^ entry (ROCE/SOCE) mechanisms[5, 6]. The transient receptor potential canonical (TRPC) family consists of seven members of the Ca^2+^-permeable nonselective cationic membrane channels within the TRP superfamily[7]. Other than the TRPC2 is encoded by a pseudogene in humans, the two TRPC1/4/5 and TRPC3/6/7 groups of subfamily members are largely expressed in the central nervous system (CNS), especially in the developing cerebellum and hippocampus[8–10]. Given the increasingly recognized roles of the TRPC family members in multiple age-related diseases[11], they have been widely considered as favored drug targets[12], including neurodegenerative diseases. However, their roles in chronic neurodegeneration are relatively understudied[13] compared to those in acute CNS conditions, which were based on the gene-ablated mouse models[14–16].

Among the TRPC family, TRPC6’s neuroprotective roles have been the most studied[17–19] and in several mouse models of familial AD. Overexpression of TRPC6 enhances spatial learning abilities in APP/PS1 mice partially via reducing the production and deposition of Aβ[20], but also by Aβ-impaired Ca^2+^ entry as well as synaptic dendritic morphology and function in APP-KI models[21]. Moreover, reduced TRPC6 levels were detected in AD patients’ sera[22, 23]. Even though these two closely related subfamily members TRPC3 and TRPC6 were often reported as TRPC3/6 to play similar or overlapping roles from earlier research[17], evidence is emerging more recently suggesting that TRPC3 may play a distinct role from TRPC6 in several experimental settings. For example, they display opposite expressional changes (e.g., upregulated TRPC3 and downregulated TRPC6) in rat hippocampus after seizure[24, 25], consistent with the originally identified pivotal roles of TRPC3 channels in the maintenance of cell excitability and induction of membrane depolarization[5, 26]. Indeed, TRPC3 channels reportedly regulate hippocampal excitability via modulating the Ca^2+^-dependent afterhyperpolarization[27]. Of note, overactivated TRPC3 channels were also found to cause maldeveloped cerebellar Purkinje neurons, underlying the ataxia phenotype in the moonwalk mice[28, 29]. Here, using an unbiased bioinformatic analysis of the published single-cell RNA-seq database from human AD brains[30], we discovered that the *Trpc3* gene is uniquely upregulated in excitatory neurons compared to the other members. Together with our findings that TRPC3 is also upregulated in aging mice and human AD brains, we speculated that TRPC3 likely plays a significant role in AD pathogenesis as in many other age-related diseases[31]. Supporting this hypothesis, knockdown of TRPC3 in hippocampus via siRNA was reported to enhance contextual fear memory[27], suggesting TRPC3 inhibition may be a viable therapeutic option for AD. Although a pyrazole-based compound (Pyr3) is widely regarded as a TRPC3-selective antagonist[32], the poor stability limits its use in *in vivo* studies. Therefore, we developed a pharmacological TRPC3 selective antagonist compound with more feasible CNS permeability and stability[33] to investigate the potentially important role of aberrant TRPC3 functions in AD pathogenesis in both *in vitro* and *in vivo* assays.

Over the past two decades, soluble oligomeric forms of Aβ (AβOs) have been recognized as the major culprit in targeting synaptic components[34], including Ca^2+^ disruption[35]. NMDA receptors are widely accepted as the principal membrane molecules responsible for Ca^2+^ overload that leads to neurotoxicity and neuronal death[36]. Potential contributions from the TRPC family have never been addressed in response to Aβ regarding Ca^2+^ signaling. Here, we report that AβOs induce *Trpc3* gene upregulation in cultured primary mature neurons that contribute to Ca^2+^ overload and neurotoxicity. Our data demonstrates for the first time that TRPC3 is a key ion channel playing a significant role in mediating Ca^2+^ overload in response to AβOs, including its unique role in intracellular Ca^2+^ handling. Treatment in both acute and chronic models challenged by AβOs with the selective compound we developed completely reverses synaptic dysfunction and memory deficits, identifying TRPC3 as a novel therapeutic target for treating AD.

## Results

### TRPC3 expression is aberrantly upregulated in autopsy AD brains

We initially performed bioinformatic analysis of the expressional levels of the genes encoding for the TRPC family based on published transcriptomic data collected from 161 AD patients and 186 normal control brains (cortices)[37]. We found the mean expression levels of *Trpc3* were 7.345 (SD = 0.336) in AD patients (n = 161) and 7.252 (SD = 0.202) in normal controls (n = 186), demonstrating a 6.7% increase in AD patients. A two-tailed t-test revealed that this difference achieved statistical significance with a *p*-value of 0.0017 (**Fig. 1A**). Significantly reduced genes encoding for TRPC1 and TRPC5 in AD brains were also detected (**Fig. S1A**). Next, we confirmed these RNA-seq data. Increased protein levels of TRPC3 and decreased TRPC6 were also detected in the cortical region of human AD brain samples (**Fig. 1B-C**). Further analysis of a published single-cell RNA-seq database[30] revealed that gene encoding for TRPC3 is uniquely upregulated (1.8-fold) in the excitatory neurons of AD brains among the entire TRPC family members (**Fig. 1D**). Interestingly, disruption of the excitatory neuronal activity has been shown to contribute to cognitive deficits in AD[38], further strengthening that increase in TRPC3 expression particularly in excitatory neurons could contribute to AD pathogenesis.

**Figure 1.**
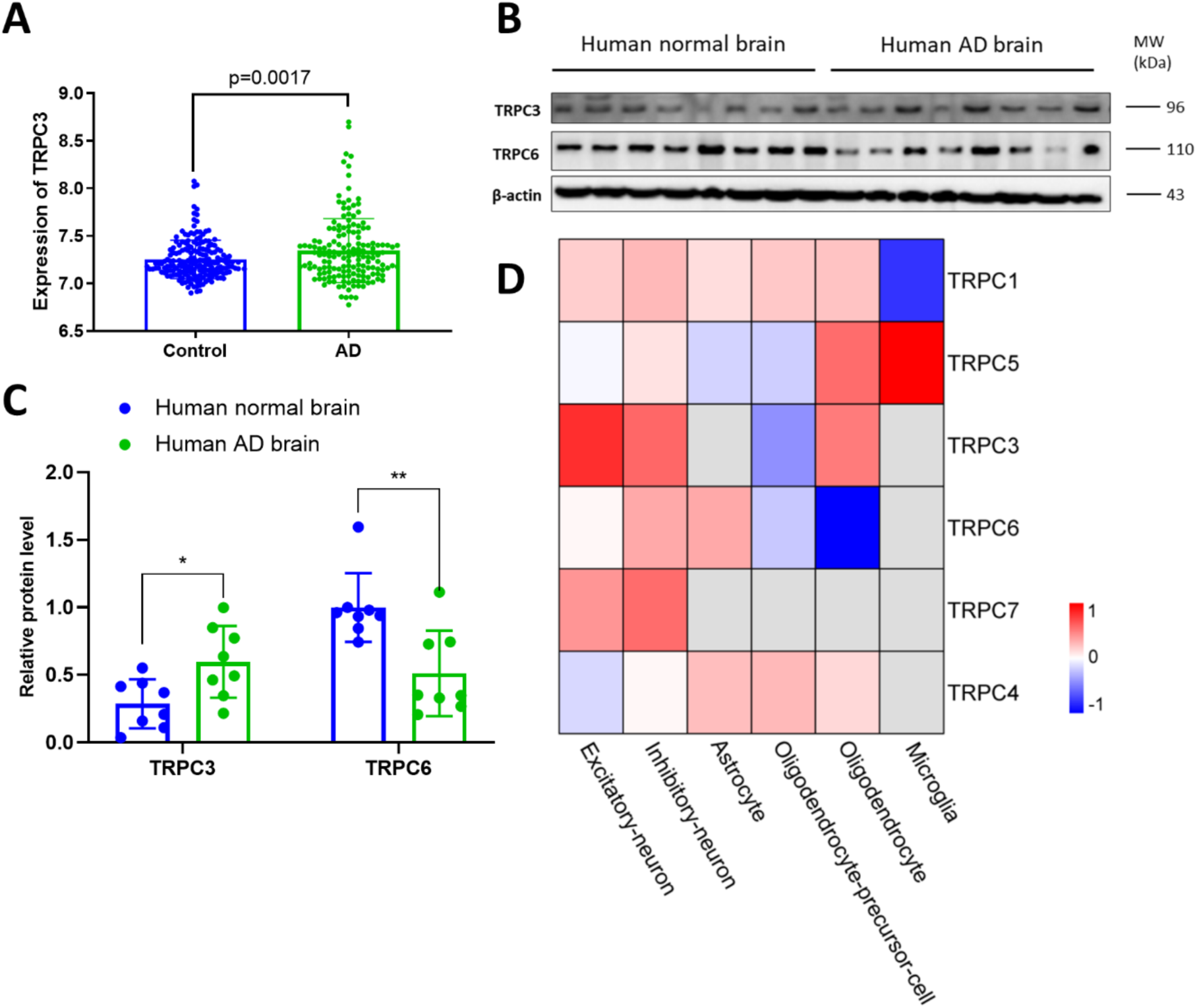
TRPC3 expression profiles in human AD samples. **A**, The expression of *Trpc3* mRNA in human patient samples[37], with n=186 for Control and n=161 for AD. **B**, The expression of TRPC3 and TRPC6 proteins in human autopsy brain. The blots are quantified in **C**. **D**, Bioinformatic analysis of TRPC family genes in AD and control brains based on single-cell RNA-seq database published[30]. Heatmap was used to show the relative expression levels between AD and control cases across different cell types. Relative expression levels were calculated using log2 Fold change between AD and control cases.

Considering age as a significant risk factor for both dementia and AD, we examined TRPC3 expression in wild-type (WT) C57BL/6 (B6) mouse brains and found that TRPC3 expression was elevated in an age-dependent manner (**Fig. S1B-C**). In addition, a familial model of AD (5xFAD Tg, which encompasses five mutations associated with AD pathogenesis) that displays features such as synaptic loss, impaired hippocampal long-term potentiation (LTP), and spatial working memory deficits as early as 4-6 months of age[39, 40], also had elevated TRPC3 levels, as compared to their littermate non-transgenic (nTg) control (**Fig. S1D-E**). On the contrary, the expressions of brain-derived neurotrophic factor (BDNF) and post-synaptic density 95 (PSD95) were reduced in the Tg samples, while TRPC6 exhibited comparable expression between the two genotypes (**Fig. S1D-E**).

### JW-65 protects against AβOs-induced damage of synaptic structure and signature proteins in vitro

The data presented above supports that TRPC3 upregulation could be associated with AD; thus, we further inhibited TRPC3 function to evaluate its role in AD pathogenesis. Using a primary culture model derived from dissected E17 embryos consisting primarily of hippocampal principal pyramidal excitatory neurons, we tested compounds that inhibit TRPC3 function. Naturally secreted oligomeric AβOs from the 7PA2 cell culture conditioned media (CM) have been well-characterized to exert synaptic toxicity in multiple *in vitro* and *in vivo* assays[41–44], and the active components were confirmed to be primarily a mixture of oligomeric forms of Aβ (i.e., monomers, dimers, trimers, and other oligomers)[45] in subnanomolar concentrations[46, 47]. Importantly, treatment of cultured mature neurons (13-14 DIV) with 7PA2 CM, but not the CM from the parental CHO cells, significantly impeded neuronal growth in a dose-dependent manner, as indicated by immunostaining of microtubule-associated protein 2 (MAP2), a neuron-specific cytoskeletal protein (**Fig. S2A**). A ten-fold diluted 7PA2 CM resulted in consistent neuronal morphological changes and cell death as we previously reported[45, 48], and thus was used throughout the subsequent experiments.

JW-65, developed by our group, is a metabolically stable, brain-penetrative, and selective TRPC3 inhibitor that can directly bind to the TRPC3 protein (**Fig. 2A**)[33]. The neuronal loss induced by 7PA2 was largely rescued by co-treatment with JW-65 (500 nM) (**Fig. 2B-C**). Furthermore, JW-65 attenuated the 7PA2-induced loss of dendritic spine structure, especially those mushroom spines generally believed to be best correlated with strong synapses and memory storage (**Fig. 2D-E**)[49–52]. Independent experiments performed on the mature primary neurons isolated from APP-KI mice[21] also demonstrated a significant rescuing effect of JW-65 on the dendritic mushroom spines (**Fig. S2B-C**). We next investigated the downstream cell signaling pathways underlying the neuroprotective effect of JW-65 by co-incubating cell cultures with a panel of kinase inhibitors commonly associated with the neuroprotective pathways. While inhibitors such as RP-cAMP (cyclic adenosine monophosphate inhibitor), LY294002 (phosphoinositide 3-kinase inhibitor), and U0126 (mitogen-activated protein kinase inhibitor) showed no significant effects, KN93, a Ca^2+^/calmodulin-dependent protein kinases inhibitor, abolished the neuroprotective effect from JW-65 (**Fig. S3A-B**); KN93 treatment alone also induced significant neurotoxicity like 7PA2 while its inactive counterpart KN92 had no effect (**Fig. S3C**). Furthermore, JW-65- treatment restored CaMKII and CaMKIV activities in specific phosphorylated forms as observed using Western blots: Thr286-CaMKII and Thr196-CaMKIV (**Fig. S3D-E**). These results suggest that the Ca^2+^/calmodulin-dependent mechanism mediates the neuroprotective action of TRPC3 antagonist against AβOs-induced synaptic toxicity. This conclusion was further supported by another signaling molecular calcineurin (CaN) downstream of calmodulin (CaM). We found a significant increase of CaN activity under 7PA2 treatment, which was rescued by JW-65 (**Fig. S3F**). Additional Western blot and RT-qPCR results also displayed a corrective effect from JW-65 on restoring levels of BDNF and PSD95, and reduced expression of the cleaved caspase-3, a marker for apoptosis (**Fig. S3G-I**).

**Figure 2.**
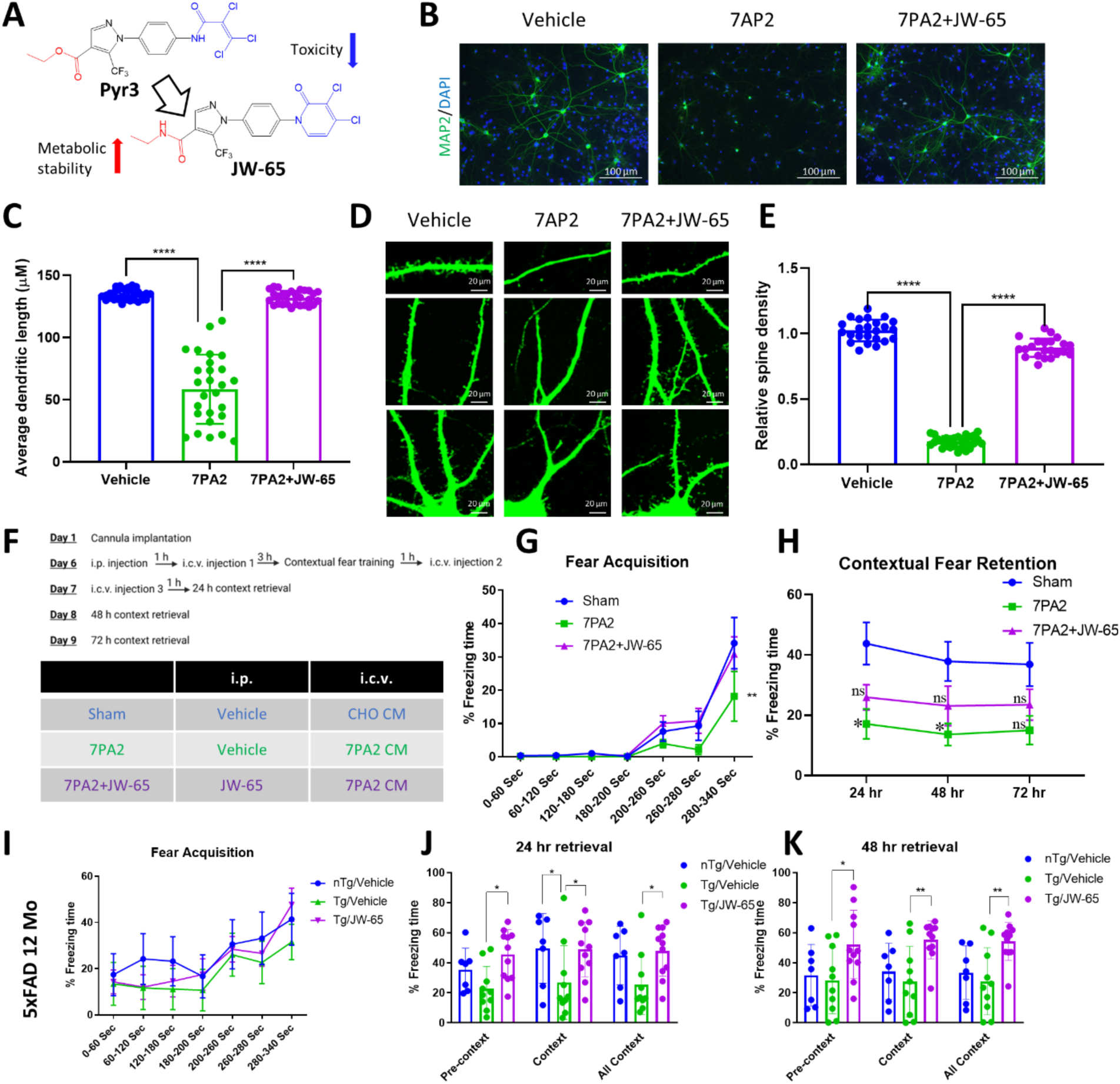
JW-65 shows neuroprotective and therapeutical effects on β-amyloidosis models both *in vitro* and *in vivo*. **A**, Chemical structure of JW-65 and its prototypic compound Pyr3[32, 33]. **B**, Representative images of cultured primary neurons treated with 7PA2 and JW-65 (500 nM). **C**, Quantification of dendritic length (n=27-30). **D**, Representative images of dendritic spine morphology under the treatment of 7PA2 and JW-65 (500 nM) by MAP2 staining. **E**, Quantification of spine density (n=22-27) as described[48]. **F**, Dosing regimen demonstrating the acute AβOs-induced synaptotoxic model. The mice received one dose of JW-65 (20 mg/kg) or vehicle post-surgical implantation through IP administration and three ICV injections of 7PA2 (2 µL) or CHO. n=4-7. **G**, Fear acquisition during the training session. **H**, Summary of the 24-72 hr retrieval test. **I-K**, Fear acquisition and 24-48 hr retrieval tests of 12-month-old 5xFAD mice. n=7- 11.

### JW-65 alleviates memory decline in both acute and chronic β-amyloidosis mouse models

We next used young adult WT B6 mice to test the optimal dose of JW65 for *in vivo* studies. Intraperitoneally (IP) injected WT B6 mice with a single dose of JW-65 up to 400 mg/Kg was well-tolerated; dosages at 20 to 100 mg/kg all led to an increase in cortical and hippocampal PSD95 protein level on Western blots, indicating that JW-65 enhanced the postsynaptic scaffolding protein expression in excitatory neurons (**Fig. S4A-B**). Further analyses using both Western blots and RT-qPCR revealed that a dosage of 20 mg/kg led to a maximum increase in hippocampal PSD95/*Dlg4* levels compared to the 10 mg/kg dosage (**Fig. S4C-D**). This dosage of 20 mg/kg was thus used in the subsequent acute and chronic *in vivo* experiments. We then tested the effect of JW-65 for hippocampus-dependent contextual memory in an acute AβOs model using 7PA2 CM as we described previously[53]. Following a single IP injection of JW-65 at 20 mg/kg, young adult WT B6 mice were intracerebroventricularly (ICV) injected with 2 µL of CHO control medium or 7PA2 CM through an implanted cannula (**Fig. 2F**). Contextual fear conditioning, a well-established paradigm for studying hippocampus-mediated context-dependent behavior facilitated by learning and memory processes, was then employed. Both context training and retrieval tests indicated that acute AβOs exposure impaired contextual learning and memory, as assessed by fear acquisition and retention processes. Intriguingly, JW-65 administration prevented this memory impairment (**Fig. 2G-H**), suggesting that inhibition of TRPC3 improves cognitive and memory functions.

To further evaluate *the in vivo* effect of JW-65, we investigated the therapeutic potential of chronic JW-65 treatment in the 5xFAD model. Initially, we characterized the phenotype of the Tg mice using a contextual fear conditioning test. Comparative analysis with nTg littermates revealed that, during the retrieval test, Tg mice displayed impaired fear contextual memory as early as five months old (**Fig. S5A-F**). This finding aligns with a prior report indicating impaired LTP in these mice starting at four months old[54]. Subsequently, we administered symptomatic Tg mice (12- month-old) with 20 mg/kg IP twice weekly JW-65 over one month. Following this therapeutic intervention, fear conditioning was conducted to assess the efficacy of the treatment. The results demonstrated that Tg mice exhibited the expected phenotype of memory decline compared to their nTg littermates. However, this memory impairment was ameliorated following JW-65 treatment (**Fig. S2I-K)**. Interestingly, bioinformatic gene analysis in the hippocampus relative to contextual fear memory across 22 strains of genetically diverse inbred BXD mice[55] revealed *Trpc3* being the sole member within the TRPC family that displayed an inverse correlation (**Fig. S5G;** *p*=0.00862), supporting our finding that TRPC3 negatively regulates contextual memory.

### JW-65 improves impaired spatial cognition and corrects dysregulated proteins in 5xFAD mice

To further evaluate the therapeutic value of JW-65, we chronically administered the compound to a new cohort of symptomatic 5xFAD Tg and nTg littermate mice, using the same treatment regimen as previously described. We specifically selected symptomatic five-month-old 5xFAD Tg mice to initiate treatment to ensure that the mice were not too advanced in age: after three months of treatment the mice could still be capable of swimming during the Morris water maze - a widely utilized behavioral test for evaluating spatial and related forms of learning and memory in rodents (**Fig. 3A**). Our results showed that, during the two training sessions, mice from the untreated Tg group were unable to reach the platform or spent the longest average time in the water before reaching the destination (**Fig. 3B-C**), indicating deficits in spatial learning and recognition. In contrast, JW-65 treatment of the Tg mice significantly mitigated their memory loss, as shown by the comparable times to reach the platform observed between them and the nTg mice (**Fig. 3B-C**). In the probe test, all groups exhibited similar swimming distances, suggesting comparable locomotor activity. However, the Tg group displayed fewer entries and spent less time in the target zone. Tg mice treated with JW-65 demonstrated enhanced awareness of the correct platform location within the pool, with both time and entries comparable to the two nTg groups (**Fig. 3D-F**), suggesting that memory is rescued by TRPC3 inhibition.

**Figure 3.**
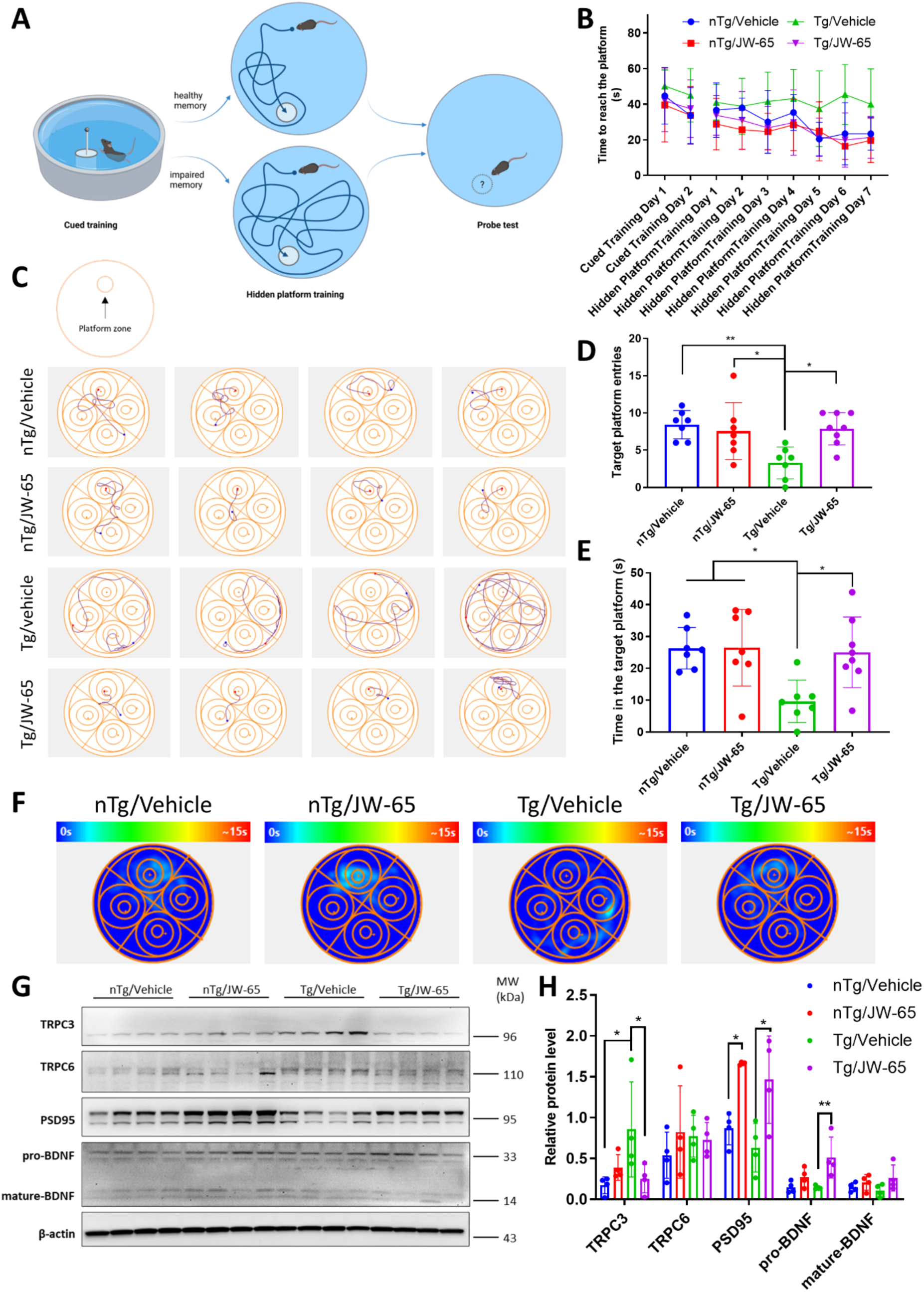
JW-65 treatment rescues spatial working memory in symptomatic 5xFAD mice. **A**, Pictorial illustration of the principle of Morris water maze. Water maze test was performed on mice at 8.0 months of age, with the JW-65 treatment group starting from 5 months of age. **B**, The time the mice spent reaching the platform during the cued training and hidden platform training sessions. n=7-8. **C**, Representative swimming traces on the hidden platform training day 7. The platform was fixed in the upper quadrant. **D-E**, Target platform zone entries and the time spent in the zone during the probe test. **F**, Representative heat maps illustrating the time spent in different quadrants during the probe test. **G-H**, Hippocampal protein expression levels of TRPC3, TRPC6, PSD95, and BDNF.

Following the completion of the water maze, the mice were euthanized, and their hippocampal protein expression was examined. Notably, Tg control mice exhibited a markedly elevated expression of TRPC3 compared to their nTg littermates. Conversely, both PSD95 and BDNF showed a significant reduction in the Tg control group (**Fig. 3G-H**). Of note, all these changes were corrected by the JW-65 treatment. On the contrary, the TRPC6 level showed no significant alterations across all experimental groups (**Fig. 3G-H**). Consequently, we infer that JW-65 effectively inhibited and decreased the hippocampal TRPC3 level and concurrently enhanced the expression of neuroprotective proteins in the 5xFAD model.

### JW-65 restores impaired LTP and rescues synaptic gene expression in 5xFAD mice

The data presented above strongly indicated that upregulated TRPC3 contributes to synaptic and memory dysfunction, which can be reversed by JW-65. Thus, we next focused on whether protection by JW-65 is mediated through synaptic function. Indeed, JW-65 was also found to restore the active synaptic zones/clusters impaired by AβOs as detected by the colocalized distribution of pre-synaptic synapsin I (SynI) and post-synaptic PSD95 as we previously described[48, 53]. Quantification based on the confocal images of dendritic spine structures with SynI-PSD95 clusters showed a significant protective effect from JW-65 (**Fig. 4A-C**). We then assessed the potential effect of JW-65 on rescuing the impacted LTP in symptomatic 5xFAD transgenic mice. LTP is a paradigm directly associated with synaptic plasticity and learning/memory. *Ex vivo* LTP recordings were conducted on brain slices from middle-aged 5xFAD mice. Field excitatory postsynaptic potential (fEPSP) was evoked in the proximal apical dendrites in hippocampal CA1b during the stimulation of Schafer-commissural projections in CA1a, and LTP was induced by theta burst stimulation (TBS) (**Fig. 4D**). Our observations revealed a significant reduction in TBS-induced LTP in old transgenic mice, suggesting an impacted LTP in the Tg mice. Conversely, the JW-65 treatment completely normalized the impaired LTP, with the fEPSP level being restored to a comparable level as observed in nTg control mice (**Fig. 4E-F**), suggesting a complete recovery of the impaired LTP. Interestingly, chronic JW-65 treatment did not seem to exert significant effects on lowering amyloid load and reducing gliosis in mid-aged mice, despite a notable impact on restoring neuronal loss and dendritic spines, as indicated by Golgi staining (**Fig. S6**).

**Figure 4.**
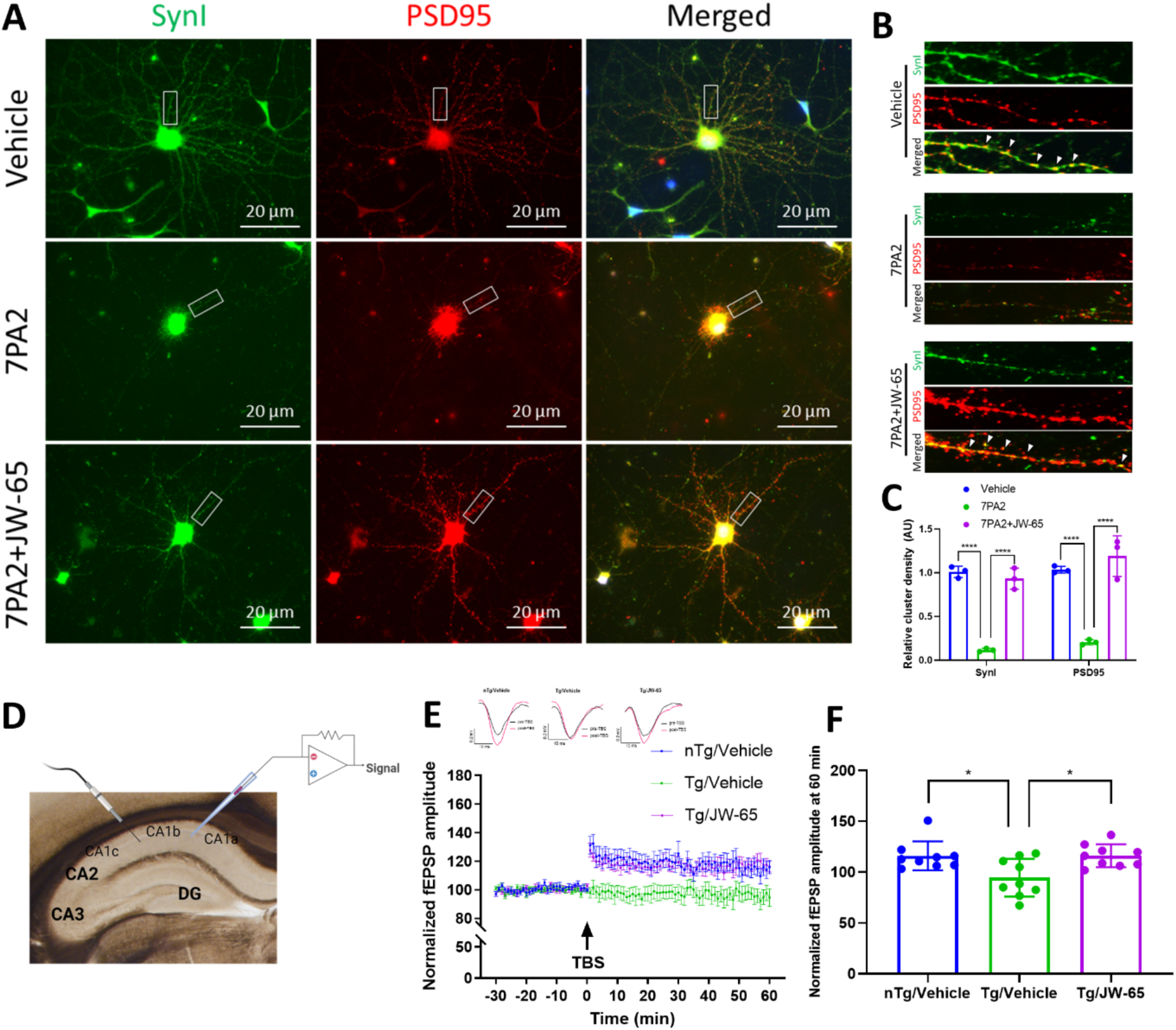
**JW-65 improves synaptic performance *in vitro* and *ex vivo*. A-C**, Confocal images of synaptic clusters based on colocalized immunosignals of Synapsin I (SynI) and PSD95. Quantification was performed as described[48]. **D-F**, *Ex vivo* LTP recording on 5xFAD brain slices. **D**, Pictorial illustration of the *ex vivo* LTP recording configuration. **E**, Time course for theta-burst induced (black arrow) LTP depicting normalized fEPSP amplitude. n=9. Top panel shows representative field synaptic responses collected during baseline (black line) and after theta burst stimulation (red line). Scale: 0.2mV/10ms. **F**, Quantification of normalized fEPSP at 60 min post-TBS. n=9.

To further uncover the molecular mechanisms underlying the synaptic protective effect of JW-65 and TRPC3 inhibition, we conducted RNA-seq experiments to capture global gene expression patterns using hippocampal total RNA from middle-aged mice. Principal component analysis (PCA) demonstrated a clear separation among the three groups, with a major variance (PC1 = 52% variance) between the nTg group and the two Tg groups (**Fig. 5A**). Further analysis revealed that 2565 genes exhibited significant differences between the Tg and nTg control littermates (*p.adj* < 0.05 and Log2 Fold Change (LFC) > 0.58) (**Fig. 5B**). Gene ontology (GO) analysis was employed to gain insights into the functional implications of the identified differentially expressed genes (DEGs). The GO analysis of BP highlighted major and significant enrichment of genes associated with synapse organization in Tg mice and immune response, indicating a prominent feature of neuroinflammation in the 5xFAD model (**Fig. 5D-E**). Tg mice in the JW-65 treatment group displayed 1293 differentially expressed genes compared to Tg control mice (**Fig. 5C**). The 994 overlapped DEGs were identified between the two compared groups, and most of those DEGs were reversed by JW-65 compared to Tg mice, including synapse organization genes like *Bdnf* as well as genes associated with Ca^2+^ ion membrane transporter activities (**Fig. 5F-H** and **Fig. S7A**). These findings are consistent with the transcriptomic data from human AD brains showing synaptic genes are largely downregulated[56–59]. Interestingly, the several major categories of microglia-associated genes including the disease-associated microglial DAM or microglial neurodegenerative phenotypes (MGnD) genes (*ApoE*, *Cst7*, *Clec7a*, Ccl2, *Ccl6*, *Trem2*, *Tgf1b, Gpnmb, Itgax, Csf1,* and *Spp1*), and the homeostatic microglial genes (*Aif1* and *Tmem119*)[60, 61] were all significantly increased in Tg mice. However, none was significantly affected by JW-65 (**Fig. S7B-D**). It is worth noting that the expression of the *Trpc3* gene was slightly increased (LFC = 0.473 < 0.58) and significantly decreased with JW-65 treatment (LFC = -0.776, *p.adj* = 9.96E-3) (**Fig. S7A**). On the contrary, *Trpc4* and *Trpc5* genes were significantly decreased in Tg but restored by JW-65 (**Fig. S7A**). Of note, reduced expression of TRPC4/5 have been shown to be associated with impaired synaptic plasticity and spatial working memory formation[62], though it is unclear how JW-65 can restore their gene expression, presumably via corrected Ca^2+^ signaling. Taken together, these findings strongly support the synaptic protection being the primary mechanism mediating the *in vivo* efficacy of JW-65 in the 5xFAD model.

**Figure 5.**
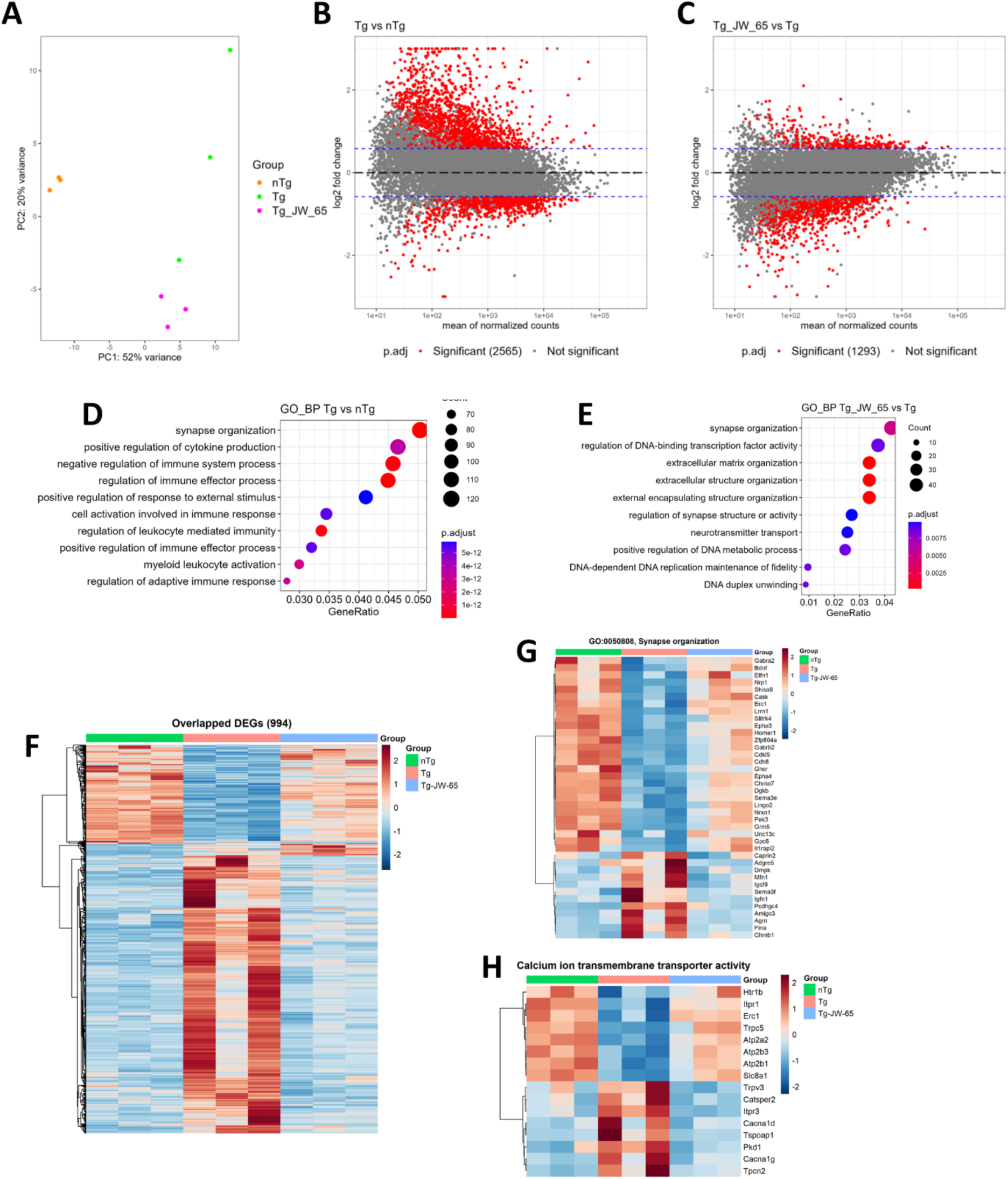
RNA-seq data analysis. **A**, PCA plot of the RNA-seq data from hippocampal tissue isolated from the brains of 5xFAD mice. nTg, n=3. Tg, n=3. Tg/JW-65, n=3. **B**, MA plot of the comparison between Tg mice and nTg mice. Red dots on the graph represent statistically significant data points. *p.adj*<0.05 and LFC>0.58. **C**, MA plot of the comparison between Tg mice received JW-65 treatment and Tg control mice. Red dots on the graph represent statistically significant data points. *p.adj*<0.05 and LFC>0.58. **D-E**, GO enrichment column graph showing the number of genes that are significantly different in Tg mice compared with nTg mice and untreated Tg compared to JW-65-treated Tg regarding the biological process (BP), related GO terms. **F-H,** Heatmaps showing the critical genes altered (*p.adj*<0.05 and LFC>0.58) by JW-65 treatment in Tg mice.

### AβOs aberrantly upregulate TRPC3 and downregulate TRPC6 expression

Since upregulated TRPC3 expression was detected in both human AD and 5xFAD Tg mouse brain tissues, we then sought to investigate if AβOs can induce this change. Indeed, 7PA2 treatment of cultured neurons rapidly induced upregulation of TRPC3 protein levels, as detected by immunocytochemistry, which can be prevented by JW-65. Time course study indicated significantly upregulated TRPC3 after 7PA2 treatment, and remained at high levels until at least 8 hr, and started declining to the basal levels at 16 hr (*Liao, unpublished data*). On the contrary, 7PA2 treatment resulted in a reduced expression level of TRPC6, a closely related TRPC member of TRPC3 (**Fig. S8A-C**). We also performed immunocytochemistry in the absence of Triton X- 100 to permeabilize the cell membrane, which allowed the detection of antigens on the cell surface. We observed the same results (**Fig. S8D-F**), indicating that AβOs induced increased expression/distribution of TRPC3 while reducing TRPC6 levels on the plasma membrane. Confocal images also display opposite changes on TRPC3 and TRPC6 after 7PA2 treatment, suggesting a reciprocal effect between TRPC3 and TRPC6 (**Fig. S8G-H**).

### TRPC3 contributes to Ca^2+^ overload during AβOs-induced neurotoxicity

In healthy neurons, cytosolic Ca^2+^ concentration is typically maintained at a lower level compared to the extracellular space or certain intracellular compartments. However, in AD, the presence of Aβ disrupts Ca^2+^ signaling by inducing an increase in intracellular Ca^2+^ levels, known as Ca^2+^ overload[63]. We conducted calcium imaging on primary cultured neurons at DIV12-14 to visualize the dynamics of Ca^2+^. In the presence of extracellular Ca^2+^, 7PA2 induced an increase in cytosolic Ca^2+^ level (**Fig. 6A**). Perfusion of 2 µM ionomycin at the end of the recording was used to achieve maximum Ca^2+^ response for inter-experiment normalization. Notably, co-perfusion of JW-65 (500 nM) with 7PA2 significantly alleviated Ca^2+^ overload (**Fig. 6A-B**). Moreover, when cells were recorded in a divalent-free (DVF) extracellular environment, 7PA2 could still increase the cytosolic Ca^2+^ level, albeit the increase was significantly lower than the increase observed when Ca^2+^ was supplemented (**Fig. 6C-D**). This suggests that 7PA2 can induce both Ca^2+^ store-release and influx, ultimately resulting in Ca^2+^ overload. In addition, JW-65 challenged both Ca^2+^ store-release and influx induced by 7PA2, as indicated by the lower fluorescence readout in both solutions (**Fig. 6C-D**).

**Figure 6.**
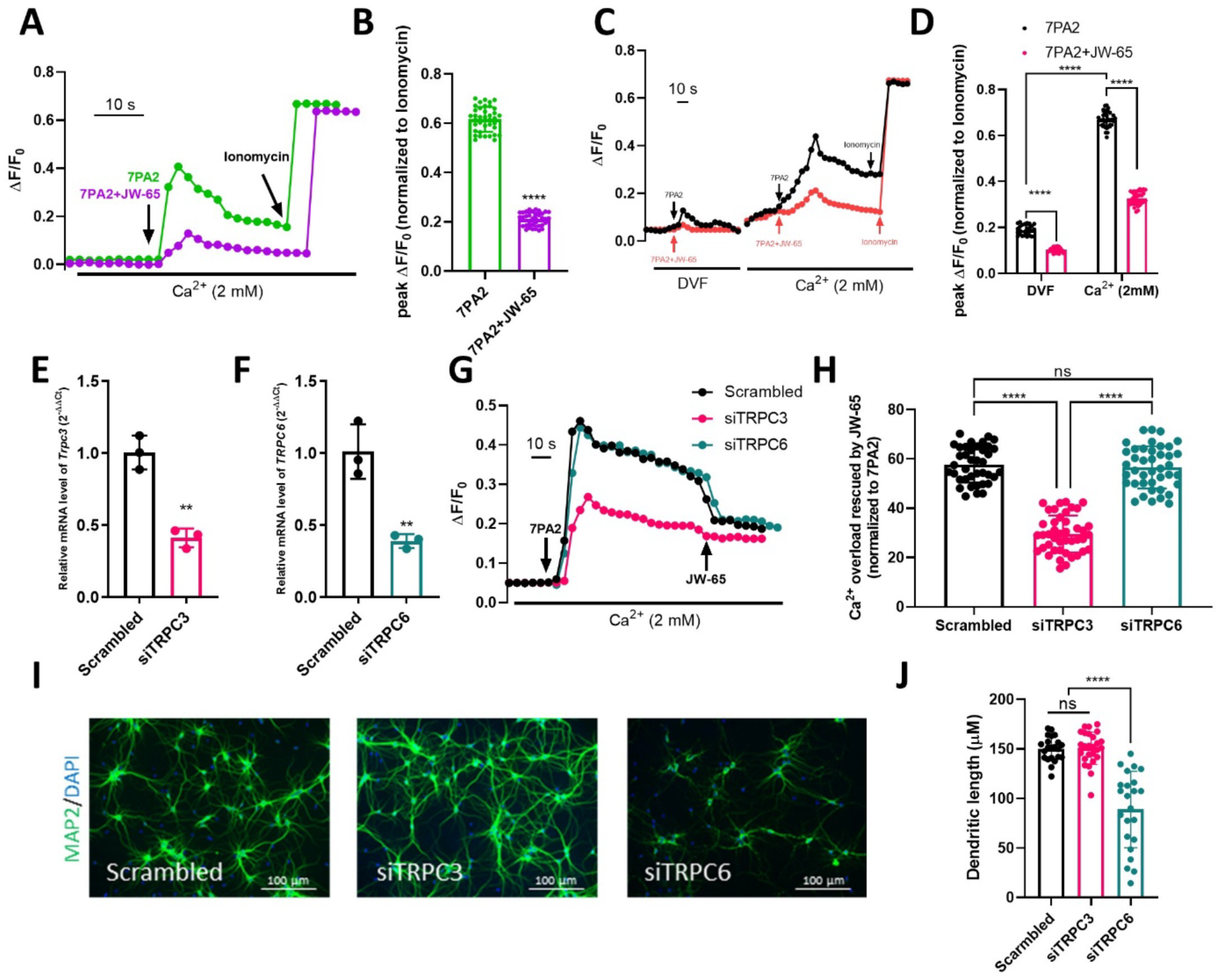
TRPC3 contributes to AβOs-induced neuronal Ca^2+^ overload. **A**, Representative fluorescent recordings on neurons perfused with 7PA2 or in combination with 500 nM JW-65, with the presence of extracellular Ca^2+^. **B**, Quantification of the peak cytosolic Ca^2+^level induced by 7PA2, normalized to ionomycin-induced overload. n=35-38. **C**, Representative fluorescent recordings on neurons perfused with 7PA2 or in combination with 500 nM JW-65 within a divalent-free (DVF) environment. The cells were then washed and exposed to an extracellular solution containing Ca^2+^ and perfused again with 7PA2 or in combination with 500 nM JW-65. **D**, Quantification of the peak cytosolic Ca^2+^ level induced by 7PA2, normalized to ionomycin-induced overload. n=27-32. **E-F**, RT-qPCR validation of the knock-down of TRPC3 and TRPC6. **G**, Representative fluorescent recordings on scrambled control, siTRPC3, and siTRPC6 neurons, with the presence of extracellular Ca^2+^. 7PA2 was perfused first to induce Ca^2+^ overload. JW-65 was then perfused to study its effects. **H**, Quantification of the Ca^2+^ overload rescued by JW-65, normalized to 7PA2-induced overload. n=39-43. **I**, Representative images of DIV14 neurons transfected with scrambled RNA, or siRNAs of TRPC3 or TRPC6. **J**, Quantification of the dendritic length in panel **I**, n = 22-26.

To confirm whether the effect of JW-65 is mediated through TRPC3, we transfected rat primary neurons with siRNAs targeting TRPC3 (siTRPC3) or TRPC6 (siTRPC6), respectively, 48 hours prior to recording. Successful knockdown was validated by RT-qPCR (**Fig. 6E-F**). In all cases, 7PA2 elevated intracellular Ca^2+^ concentration. However, the elevation of Ca^2+^ was significantly lower when TRPC3 was knocked down (**Fig. 6G-H**). Importantly, JW-65 only exerted a minor effect when the expression of TRPC3, but not TRPC6, was knocked down (**Fig. 6G-H**), indicating TRPC3 as its bona-fide target. While it is known that Aβ induces Ca^2+^ overload by interacting with various channels, including ligand-gated calcium channels such as NMDA receptors (NMDARs), plasma membrane Ca^2+^ ATPase (PMCA), sodium-calcium exchanger (NCX), voltage-gated calcium channels (VGCCs), as well as inositol 1,4,5-triphosphate receptors (InsP3Rs) and ryanodine receptors (RyRs) on the membrane of endoplasmic reticulum (ER)[64], our results suggest that TRPC3, as a channel protein, is also significantly involved in this process. Relative contributions of TRPC3 versus these channels to the AβOs-induced Ca^2+^ overload in neurons, and the potential interplay between TRPC3 with some of them (e.g. NMDARs and VGCCs) warrants further investigation. In addition, electroporation was performed to silence TRPC3 and TRPC6 on DIV0. Surprisingly, on DIV14, a stage of neuronal maturity, a dramatic difference was observed between the effects of knocking down TRPC3 and TRPC6. While TRPC6 appeared to be highly neuroprotective, TRPC3 showed minimal impact on neuronal morphology. This suggests distinct functions between the two close homologs (**Fig. 6I-J**).

## Discussion

### Aberrantly upregulated Trpc3 in glutamatergic excitatory neurons

We report here that TRPC3 expression is aberrantly upregulated during AD progression (human and mice models), correlating with impaired synaptic structures, plasticity, and fear contextual as well as long-term spatial working memory in multiple *in vitro* and *in vivo* experimental paradigms. Application of a newly developed pharmacological antagonist compound JW-65 restores TRPC3 expression levels and completely corrects all these synaptic dysfunctions. A further mechanistic investigation based on soluble Aβ oligomers as the major AD culprit, we found that AβOs can induce TRPC3 upregulation in neurons, contributing to Ca^2+^ overload, dendritic damage, and neuronal death, which again can be completely prevented/rescued by TRPC3 inhibition via JW- 65 treatment. Given the well-recognized concept that AD is a fundamental disease of synaptic failure[65, 66], the several findings disclosed in this work not only identify a promising pharmacological agent with translational potential but also pinpoint several mechanistic insights of TRPC3 uniquely contributing to the pathogenesis and progression of AD. Our bulk hippocampal RNA-seq data strongly indicates that the upregulation of the *Trpc3* gene contributes to synaptic failure and impaired learning/memory, and thus plays a significant role in AD pathogenesis and progression. Furthermore, we find that AβOs can induce TRPC3 upregulation in excitatory neurons at the transcriptional level as a result of the activated Ca^2+^/CaN-NFAT downstream of Ca^2+^ overload (*Liao, unpublished data*). Interestingly, our bulk hippocampal RNA-seq analysis also identifies multiple genes related to Ca^2+^ homeostasis/signaling that are altered in symptomatic 5xFAD mice but are largely corrected by TRPC3 blockage via JW65. We therefore speculate that JW65’s effect to correct upregulated neuronal expression of TRPC3 is primarily owing to its ability to restore Ca homeostasis after AβOs challenge, and thus correcting the overactivated Ca^2+^/CaN-NFAT signaling, resulting in normalized *Trpc3* gene at transcriptional level.

### How does the upregulated Trpc3 expression in excitatory neurons contribute to synaptic failure and AD pathogenesis?

Through testing the selective TRPC3 antagonist compound in the chronic 5xFAD mouse model, we learned that aberrantly upregulated TRPC3 negatively correlates with impaired synaptic morphology and functions. Moreover, results obtained from our *ex vivo* AβOs-challenged neuronal culture model and hippocampal slice recording corroborate well, demonstrating that TRPC3 overactivation can contribute to AD synaptic failure in conjunction with amyloid pathology. Interestingly this forms a vicious cycle, where AβOs induce TRPC3 upregulation, as a result of Ca^2+^ overload, which contributes to sustained Ca^2+^ dyshomeostasis. We demonstrate that JW-65 treatment can rescue impaired LTP in the hippocampus of 5xFAD Tg mice, which is consistent with an earlier report based on a mouse genetic approach from crossing TRPC3 global knock-out (KO) mice with the 5xFAD, showing corrected fear contextual memory[27]. The same group also reported decreased basal neuronal excitability and an increase in Ca^2+^-sensitive spike afterhyperpolarization (AHP) detected in a subset of 5xFAD weak-learner mice[67]. These results suggest a potential function of the elevated TRPC3 expression and overactivation in contributing to impaired LTP through promoting the AHP.

In parallel, our data also indicate that overactivated TRPC3 can contribute to excitotoxicity presumably through dysregulated intracellular Ca^2+^. Our findings identify TRPC3 from the TRPC family as a key ion channel, alongside the classically defined NMDARs and VGCCs, causing Ca^2+^ overload upon Aβ stimulation. On the other hand, our recent report demonstrates a potent anti-seizure effect of the same TRPC3 antagonist compound JW-65 in mice[68], strongly arguing a significant role of overactivated TRPC3 in hyperexcitability, an increasingly reported phenomenon in the early stage of AD[69, 70]. TRPC3 is reportedly upregulated during seizure in rodent models[24, 25]. Although seizures were not detected in 5xFAD mice at all ages despite the detected hyperactivity, the most notable behavioral phenotypes of the 5xFAD model[71], surprisingly, hyperexcitability was reported in 5xFAD at old age in one study[72]. On the contrary, an earlier report suggests insufficient learning-related intrinsic excitability (hypoexcitability) underlies synaptic failure in middle-aged 5xFAD mice[67]. Our data gathered from the studies with JW-65 in middle-aged 5xFAD mice are more in line with the notion that overactivated TRPC3 contributes to impaired LTP or hypoexcitability. Future work should clarify the seemingly contradictory roles of overexpressed/overactivated TRPC3 in hyperexcitability versus hypoexcitability during AD progression, both are present in different stages of AD patients[69, 70].

### TRPC3-BDNF axis in the context of synaptic plasticity in AD

Given the prominent roles of BDNF in regulating synaptic plasticity and the cognitive functions of the brain, we reason that JW-65 rescues hippocampal LTP at least partially via restoring BDNF functions. Indeed, the JW-65 treatment restored *Bdnf* gene transcription in cultured neurons and 5xFAD hippocampal tissues. Furthermore, JW-65 treatment also consistently restores AβOs-impaired *Dlg4/*PSD95, likely as a result of resumed Ca^2+^ homeostasis via Ca^2+^/calmodulin-mediated synaptic protective mechanisms involving CaMKII/CaMKIV activation[73, 74]. Despite high amyloid plaque load and substantial neuroinflammation, the treated 5xFAD mice display normal learning and memory. Again, this attests that AD is fundamentally a disease of synaptic failure, as evidenced by the impaired LTP in the hippocampus of the 5xFAD mice. Interestingly, all excitatory neuron subtypes express high levels of BDNF. Although impaired BDNF signaling has been linked with increased amyloid and tau pathology, neuroinflammation, and neuronal loss, the exact mechanisms underlying the effect of impaired BDNF signaling on AD are still largely unknown[75]. BDNF signaling through its major receptor TrkB requires Ca^2+^ influx and the activation of Ca^2+^/CaMKII[76]. Therefore, our data support a theory that defective BDNF signaling in AD may be a result of dysregulated Ca^2+^ and the Ca^2+^/calmodulin-dependent signal transduction in which TRPC3 overactivation may play a part. BDNF-induced tropomyosin receptor kinase B (TrkB) activation mediates PLC-dependent TRPC3 channel activity and Ca^2+^ entry in pontine neurons during development[77]. In addition, TRPC3 channel activity is essential for nerve-growth-cone guidance by BDNF in cerebellar granule cells[78]. Of particular interest, TRPC3 was found to be required for the BDNF-TrkB-induced slow depolarization in CA1 pyramidal neurons to promote dendritic spine density[79]. Moreover, BDNF from mossy fibers evokes a TRPC3 current and Ca^2+^ elevations in CA3 pyramidal neurons[80]. All these prior reports identify TRPC3 as the mediator of BDNF-induced membrane current channel imperative for the BDNF signaling via TrkB under physiological conditions. On the contrary, our work discloses a novel association between aberrantly upregulated TRPC3 and impaired BDNF signaling during AD progression, both are attributed to calcium overload, which potentially involves overactivated TRPC3 channel activity to be determined.

### Distinct roles of TRPC3 and TRPC6 during AD progression

In earlier reports, TRPC3/6 were often reported together for overlapping roles in physiological functions, such as promoting cerebellar granule neuron survival[17]. Although both of them are highly expressed in CNS in terms of the hippocampus, they appear to be expressed in different hippocampal subregions[8–10, 81], implying that they may play distinct roles in various neuronal subtypes. TRPC6 has been found crucial in the formation of excitatory synapses[82]; however, the scRNA-seq data from AD patients indicates that TRPC6 expression in excitatory neurons displays no changes compared to markedly elevated levels of TRPC3 (**Fig. 1D**). In fact, other than TRPC3, genes encoding for the other members were all found expressed in GABAergic inhibitory neurons in the CA3 of the hippocampus[83], in various combinations. TRPC3/6 frequently form in homomers while TRPC1/4/5 are in various heteromers[84]. It remains to be determined the relative or different roles of TRPC3 and TRPC6 in excitatory as well as in inhibitory neurons during AD progression in terms of hippocampal formation[85]. In fact, TRPC3 and 6 heterodimers have never been identified thus far under physiological conditions. They may be expressed in different neuronal subpopulations. More recently, divergent roles of TRPC3 and TRPC6 have been increasingly reported especially in neuronal hyperexcitability and epilepsy[24, 25]. Seizure induces upregulated TRPC3 and downregulated TRPC6, respectively[24]. TRPC6 has also been shown to be neuroprotective, and pharmacological TRPC6 agonists such as hyperforin and piperazine derivatives have also been widely reported to be promising agents for treating AD[21, 86, 87]. However, it remains to be determined how TRPC3 and TRPC6 are differentially regulated by AβOs. Our data show that TRPC3 upregulation is at the transcriptional levels via activated CaN/NFAT as an event downstream of Ca^2+^ overload. On the other hand, TRPC6 downregulation appears to be primarily at protein levels via lysosome-mediated degradation (**Supporting File**). Differential distributions of these two members in different intracellular compartments, such as TRPC3 in mitochondria[88] and TRPC6 in ER[20] suggest their potentially distinct roles in intracellular Ca^2+^ handling, which warrants further investigation. As the TRPC family is increasingly becoming a favored class of therapeutic targets for age-related diseases such as AD, a fuller understanding of the potentially distinct roles of each TRPC member plays during disease onset and progression is imperative and instrumental for rationalizing drug targeting strategies and treatment options.

### Neuronal calcium signaling as a therapeutic target for AD

There is extensive previous literature suggesting that dysregulated Ca^2+^ signaling plays a key role in AD pathogenesis and that Ca^2+^ signaling modulators may have potential neuroprotective effects in AD[89, 90]. The targets that were previously considered in this context include RyR2[91] and our recent genetic studies demonstrated that increased basal activity of RyR2 inhibits autophagy in AD neurons[92]. Another potential target is CaN. Retrospective analysis of epidemiological data revealed a significantly reduced incidence of AD in transplant patients chronically treated with CaN inhibitor FK-506[93]. A recent report based on a large, diversified patient population with controls for age, sex, race, and multiple AD risk factors disclosed that the use of the FDA-approved CaN inhibitor, tacrolimus is also associated with reduced prevalence of dementia-related symptoms[94]. Our recent results suggested that positive allosteric modulators of SERCA Ca^2+^ pump (NDC compounds) may also stabilize neuronal Ca^2+^ signaling and exert neuroprotective effects in AD models[95, 96]. In the present study, we report that similar synaptoprotective and neuroprotective effects can be achieved by selective TRPC3 inhibitor JW-65.

As illustrated in Graphical abstract (**Fig. S9**) exemplified by AβOs as a challenge in AD neurons, inhibitors of TRPC3 and other Ca^2+^ signaling modulators may act by preventing calcium overload via suppressing the overactivation of CaN as well as preserving the calmodulin-dependent kinases CaMKs, resulting in synaptoprotection. Because of these convergent mechanisms of action, the aforementioned compounds may also exert synergistic effects when used in combination. Ultimately, the choice of Ca^2+^ signaling modulators for future therapeutic development will be determined by their relative efficacy and safety profiles.

## Materials and methods

### Study approval

All animal procedures were performed in accordance with the Animal Scientific Procedures Act and with the approval of the Institutional Animal Care and Use Committee (IACUC) at the University of Tennessee Health Science Center. The 5xFAD breeder mice were originally purchased from JAX (Stock #034840) and were bred via mating hemizygous transgenic males with wild-type B6SJLF1 breeders (Jackson Labs, Stock #100012), with the mutant retinal degeneration allele (Pde6b+/rd) bred out of the colony.

### Human transcriptome analysis

The single-cell transcriptomics data published by Mathys et al[30] were collected from 24 individuals with AD pathology and 24 healthy controls, which contains 6 major cell types, including excitatory neurons, inhibitory neurons, astrocytes, oligodendrocytes, oligodendrocyte precursors, and microglia cells. We extracted 6 TRPC family members (TRPC1, TRPC3, TRPC4, TRPC5, TRPC6, and TRPC7) across various cell types.

### Hippocampal transcriptome profiling of BXD mice

The total hippocampal RNA from 22 BxD mice strains at 2∼3-month-old was used in gene array using Illumina Sentrix Mouse-6.1 bead array (http://www.illumina.com/). Raw microarray data were normalized using the rank invariant method and background subtraction protocols provided by Illumina as part of the BeadStation software suite. We log-transformed the expression values and stabilized the variance of each array to a mean of 8 and an SD of 2 per array. A single unit on this scale (log2) corresponds to a two-fold difference in expression.

### Primary neuronal culture and transfection

Primary cortical neurons were isolated from E17 embryos of Sprague Dawley rats and cultured as described previously[45, 48]. A pregnant female rat was sacrificed on E17. The embryos were obtained from oviduct and was immediately placed in ice-cold dissection solution, which is HBSS (Thermo Fisher Scientific, 14025076) medium containing 4 µL/mL glucose (Thermo Fisher Scientific, A2494001), 5 mM HEPES (Thermo Fisher Scientific, 15630130) and 1xPen/Strep (Sigma-Aldrich, P4333). The hippocampus was isolated and digested with TrypLE (Thermo Fisher Scientific, 12604013). The cells were then seeded to 100 mm cell culture dishes (Corning, 430293), for Western blot and RT-qPCR, or 24-well plates (Corning, 3524) coated with coverslips (Neuvitro, GG-12) pre-treated with poly-D-lysine (Thermo Fisher Scientific, A3890401), for staining and calcium imaging. The digested cells were seeded with plating medium, which is DMEM medium (Thermo Fisher Scientific, 11965092) that contains (per 500 mL DMEM) 62.5 mL fetal bovine serum (Cytiva HyClone, SH30071.03IR25), 5 mL GlutaMAX (Thermo Fisher Scientific, 35050061), 15.6 mL HEPES (Thermo Fisher Scientific, 15630130), 62.5 mL Ham’s F-12 Nutrient Mix (Thermo Fisher Scientific, 11765054), and 1x Pen/Strep (Sigma-Aldrich, P4333). Two hours later, the plating medium was replaced with culture medium, which is Neurobasal medium (Thermo Fisher Scientific, 21103049) that contains (per 500 mL Neurobasal) 10mL B-27 supplement (Thermo Fisher Scientific, A1486701), 5 mL GlutaMAX (Thermo Fisher Scientific, 35050061), and 1x Pen/Strep (Sigma-Aldrich, P4333). Fresh culture medium (1:2 replacement of starting volume) was added every 3 d. Transient transfections of primary neurons with siRNA were conducted by electroporation using Lonza Amaxa Rat Neuron Neucleofector kit (Lonza, VPG-1003) according to manufacturer’s instructions: 4-5x10^6^ cells were transfected with 30-300 nM siRNA.

### Naturally secreted AβOs-containing conditioned medium (7PA2 CM)

The 7PA2 cell line represents Chinese Hamster Ovary (CHO) cells stably transfected with human APP751 which contains a Val717Phe Mutation[41]. Conditioned medium (CM) containing naturally secreted AβOs was collected from 7PA2 cells as previously described[45]. 7PA2 CM and CHO CM were prepared by culturing 7PA2 or CHO cells. The culturing medium is DMEM (Thermo Fisher Scientific, 11965092) containing 10% FBS (Cytiva HyClone, SH30071.03IR25), 200 µg/mL G418 (Gibco, 10131035), and 1% Pen/Strep (Sigma-Aldrich, P4333). Upon 80% confluency, collecting medium that is Neurobasal medium (Thermo Fisher Scientific, 21103049) containing 2 mL (per 500 mL Neurobasal) 200 mM L-glutamine (Gibco, A2916801) and 1% Pen/strep (Sigma-Aldrich, P4333) was added to replace the original culturing medium. After 24 h, the collecting medium was filtered (Millex, SLG004SL) and collected.

### Chemical compounds and reagents

JW-65 was synthesized in-house as described[33]. KN92 (4130) and KN93 (1278) were purchased from Tocris Bioscience; RpcAMP from Sigma-Aldrich (A165); LY294002 from LC laboratories (L-7962); U0126 from Cell Signaling Technology (9903S).

### Immunoblotting, immunocytochemistry and immunohistochemistry, RNA isolation, reverse transcription, and RT-qPCR

These procedures were performed as described[45, 48, 53]. Images were acquired by Keyence microscope BZ-X800 (Keyence) or Olympus FV1000 confocal laser scanning biological microscope (Olympus Life Science). Quantification of amyloid load, neuroinflammation, and dendritic spine were described[48, 53]. Primers and antibodies used in this research are listed in **Table S1-2**.

Cells or brain tissues were homogenized in RIPA buffer (Thermo Fisher Scientific, 89901) and protease inhibitor (Roche, 46931160010) and rested for 30 min on ice. Homogenates were centrifuged at 12,000 g for 20 min at 4 °C. Supernatants were used as total protein lysates and were analyzed by SDS-PAGE (Invitrogen, XP04200BOX). EZ-Run Prestained Protein Marker was used for Western blotting (Thermo Fisher Scientific, BP-3601). The antibodies used for Western blotting were TRPC3 (Alomone Labs, ACC-016), TRPC6 (Alomone Labs, ACC-017), PSD95 (Invitrogen, 51-6900), BDNF (Alomone, ANT-010), cleaved caspase-3 (Cell signaling Technology, 9661), β-actin (Sigma-Aldrich, A2228) and horseradish peroxidase (HRP) conjugated secondary antibodies (Thermo Fisher Scientific, Goat anti-Rabbit, 31460; Goat anti-Mouse, 31430). Blots transferred onto Immobilon PVDF membranes (Millipore, IPVH00010) were visualized via Gel Doc 2000 Imaging System (Bio-Rad).

Cells (prepared on coverslips) were washed with PBS (prepared in house) and fixed with 4% PFA (prepared in house) for 20 min. The cells or permeabilized with 0.4% Triton X-100 (Fisher Scientific, BP151-100) in PBS for 5 min. The cells were then blocked with 5% Bovine Serum Albumin (Miltenyi Biotec, 130-091-376) and 0.4% Triton X-100 (Fisher Scientific, BP151- 100) in PBS for 1 h before incubation with primary antibodies at 4 °C overnight. After washed with PBS, the cells were incubated with DAPI (Invitrogen, D1306) and secondary antibodies for 2 h at room temperature. Finally, the coverslips were mounted using Fluoromount Aqueous Mounting medium (Sigma-Aldrich, F4680). Alternatively, for membrane protein visualization, Triton X-100 was excluded from all procedures.

Deeply anesthetized mice were transcardially perfused with 30 mL of filtered PBS (prepared in house). Brains were removed and fixed in 4% PFA (prepared in house) for 24 h at 4 °C. Brains were sectioned using Leica VT 1200S vibratome (Leica Biosystems). The brain slices were blocked with 5% Bovine Serum Albumin (Miltenyi Biotec, 130-091-376) and 0.4% Triton X-100 (Fisher Scientific, BP151-100) in PBS for 1 h before incubation with primary antibodies at 4 °C overnight. After washed with PBS, the slices were incubated with DAPI (Invitrogen, D1306) and secondary antibodies for 2 h at room temperature. Finally, the slices were mounted using Fluoromount Aqueous Mounting medium (Sigma-Aldrich, F4680).

Total RNA was extracted using TRIzol (Thermo Fisher Scientific, 15596026) and quantified by NanoDrop 2000c Spectrophotometer (Thermo Fisher Scientific, ND2000CLAPTOP). 500 ng of RNA was reverse transcribed into cDNA with High-Capacity cDNA Reverse Transcription Kit (Thermo Fisher Scientific, 4368814). RT-qPCR was performed on Mastercycler Realplex Real-time PCR System (Eppendorf) using SsoAdvanced Universal SYBR Green Supermix (Bio-Rad, 1725270). The reaction used white 96-well plates (Azenta, 4ti- 0741). 2-ΔΔCt calculation was performed on triplicate Ct values normalized with housekeeping gene Gapdh or bActin. Primers purchased from Invitrogen were used at a concentration of 100 nM. Images were acquired by Keyence microscope BZ-X800 (Keyence) or Olympus FV1000 confocal laser scanning biological microscope (Olympus Life Science).

### Calcineurin activity

Calcineurin activity was determined using a Calcineurin cellular activity assay kit (Enzo Lifesciences, BML-AK816-0001).

### Dendritic spine analysis in primary hippocampal neuronal cultures

The hippocampal cultures of APPKI and WT mice were established from postnatal day 0- 1 pups and maintained as described[21]. For the assessment of synapse morphology, hippocampal cultures were transfected with TD-Tomato plasmid at DIV7 using the calcium phosphate method, and the culture was treated with different concentrations of JW-65 on DIV15. The cultures were then fixed with 4% formaldehyde 24 hr later. A-Z-stack of optical section was captured using 100x objective with a confocal microscope (Carl Zeiss Axiovert 100M with LSM510). Quantitative analysis for dendritic spines from 18-20 neurons each genotype was performed with the NeuronStudio software package[97]. To classify the shape of neuronal spines in culture, we adapted an algorithm from a previously published method[97] for quantification.

### Calcium imaging

The experimental setup (Oregon Green 488 BAPTA-2/AM, Invitrogen, O6809) and the recording conditions were previously described[98]. Extracellular solution was prepared to contain (in mM) 140 NaCl, 6 KCl, 2 CaCl2, 1 MgCl2, 10 glucose, and 10 HEPES. Divalent-free solution was prepared to contain (in mM) 145 NaCl, 5 EGTA, 2 EDTA Na+ salt, 10 glucose, and 20 HEPES. Oregon Green 488 BAPTA-2/AM (Invitrogen, O6809) was diluted into the solutions at a working concentration of 4-5 µM. Cells were incubated with the above solutions for 1 h. Fluorescent imaging was performed using a Lambda 10-2 Optical Filter Changer Control System (Sutter Instrument) at a sampling rate of 500 ms, recorded by MetaFluor software (Molecular Devices).

### Human brain specimens used in Western blot analysis

AD and non-AD brain specimens were collected from the Human Brain and Spinal Fluid Resource Center, which is sponsored by NIHDS/NIMH, the National Multiple Sclerosis Society, and the Department of Veterans Affairs. Samples were collected from short-postmortem interval (< 4 hr PMI) autopsies from the University of Kentucky AD Center (UK-ADC) cohort. The average age of individuals with AD was 68 ± 8.42, and 79 ± 3.6 for healthy controls.

### Acute AβOs-induced synaptic toxicity assay

Wild-type C57BL/6 mice (2-4 months of age, both sexes) received 2 μl of control (CHO medium), or 7PA2 CM, via an implanted cannula. Surgical implantation and intracerebellar ventricular injections were performed as described[48, 53]. Mice were anesthetized by intraperitoneal injection of ketamine/xylazine (87/13 mg/kg). Analgesic was administered (carprofen, 5 mg/kg). The eyes were applied with eye lubricant and shaded with soft tissue to avoid dehydration over time and damage caused by exposure to the light. The mice were positioned in a stereotaxic mouse frame (Kopf instruments). After being precisely orientated, the skull was exposed, and a hole was drilled into the skull (coordinates to bregma: anteroposterior (AP), -0.37 mm for mice weighing 21-25 g or -0.46 mm for mice weighing 26-30 g; mediolateral (ML), 0.95 mm for mice weighing 21-25 g or 1.00 mm for mice weighing 26-30 g). A guide cannula (Plastics One) designed to reach -1.75 mm dorsoventrally (DV) was mounted to the skull. The scalp, together with the guide cannula, was fixed with superglue or dental cement. A paired dummy cannula (Plastics One) was fitted into the guide cannula to prevent clogging. 1 mL 0.9% NaCl saline was i.p. injected for rehydration. The mice were placed on a heat pad and monitored until complete recovery from anesthesia before returning to the housing cages. 5 mg/kg carprofen was administered QD for the first three days post-surgery. Upon recovery, the dummy cannula was removed and 2 µL CHO/7PA2 CM was injected via an internal cannula (Plastics One) at an injected speed of 0.4 µL/min, controlled by fixing a microinjection syringe (Hamilton) on a syringe pump (Harvard Apparatus). After injection, the internal cannula was kept in place for an additional 5 min and slowly withdrawn to avoid liquid overflow. Finally, the Dummy cannula was placed back.

### Mouse behavioral tests

Fear conditioning was conducted as described[53, 99]. Fear conditioning consists of two sessions: A training session and a retrieval session. During the training session, each mouse was placed in a chamber (Actimetrics) with a camera on the ceiling and conductive grills as the floor. They were allowed to explore the chamber for 200 s (pre-context). After that, a white noise (80 dB and 2 Hz) followed by a foot shock (0.5 mA) was given as a stimulus two times at 200 s and 280 s, respectively, to the mice (context). Once finished (6 min in total), the mice were placed back to their housing cages. After 24-72 h, the fear retrieval test was performed several times by placing the mice back to the same chamber for 6 min but without stimulus. Freezing behavior during all the sessions was recorded and analyzed by FreezeFrame 3 software (Actimetrics).

The Morris water maze testing paradigm consists of three sessions as we previously described[53]. Spatial learning and memory were tested by Morris water maze test. A circular tank (1 m in diameter and 40 cm in height) was filled with opaque white water at a depth of 20 cm. Different shape stickers were placed along the walls surrounding the tank as reference cues. An escape platform was placed at the center of one quadrant. A rod with a conspicuous plastic ball on top was fixed vertically on the platform in the cued training session, during which the platform stayed above the water surface. Mouse behavior was video recorded. During cued training, mice were introduced into the pool from four different quadrants, and were allowed to search for the platform for 60 s. If their escape failed, they were guided to the platform and kept on it for 10 s. During hidden platform training, the rod on the platform was removed and the platform was pushed down to be hidden 1 cm underneath the water surface. The same 60 s tests were performed as the cued training test. After training, a probe test was performed by removing the platform and allowing each mouse to swim freely for 60 s. The swimming traces during the three sessions were analyzed via ANY-maze (Stoelting).

### RNA-sequencing analysis

Raw RNA-sequencing analysis was processed with Salmon[100]. Reads were aligned to the Ensembl mouse reference genome GRCm39 and the gene expression was quantified. The raw gene counts were processed in RStudio (Ver2022.07.2+576) with DESeq2 package to analyze differentially expressed genes (*p.adj* < 0.05 and Log2 Fold Change (LFC) > 0.58).

### Recording of long-term potentiation (LTP) of CA1 field excitatory postsynaptic potentials (fEPSPs) in hippocampal brain slice preparations

The experimental setup (coronal brain slices at 400 µm thickness containing the hippocampus) and the recording conditions were previously described[101, 102]. Mice were deeply anesthetized with isoflurane, and brains were rapidly removed and transferred to ice-cold prep solution containing (in mM) 5 KCl, 1.25 NaH2PO4, 5 MgSO4, 1 CaCl2, 205 sucrose, 25 glucose, and 26 NaHCO3 (pH 7.4, Osm 300-310) under 95% O2 and 5% CO2. Brains were sectioned using Leica VT 1200S vibratome (Leica Biosystems) and incubated in artificial cerebrospinal fluid (ACSF) containing (in mM) 115 NaCl, 2.5 KCl, 2 CaCl2, 1 MgCl2, 1.25 NaH2PO4, 25 NaHCO3, 11 glucose, and 0.1 vitamin C (pH 7.4, Osm 280-310) under 95% O2 and 5% CO2 for 2 h at room temperature. Upon recording, a 400-µm slice was transferred into the submerged recording chamber and superfused with ACSF. The Schaffer collateral/commissural fibers in the stratum radiatum were stimulated with a bipolar tungsten electrical stimulating electrode (76 mm in length, 0.5 MΩ nominal impedance. Tungsten, TST33C05KT). Field excitatory postsynaptic potentials (fEPSPs) were recorded from the CA1 stratum radiatum using an extracellular glass pipette filled with ACSF placed 200-300 mm away from the stimulating intensity that yielded 50% of the maximum response. After a 30 min baseline recording, theta-burst stimulation was used to induce LTP, followed by continued recording for 60 min.

### Golgi staining

Golgi staining was performed as previously described[103] using the FD Rapid GolgiStain Kit (cat. # PK401, FD Neurotechnologies, Inc., Ellicott City, MD 21041, USA). Spines were quantified with Fiji software[104].

### Statistical analysis

All data are presented as mean±SD. Sample sizes are described in the figure legends. Statistical analysis was performed using GraphPad Prism (Ver 8.3.0). Two-tailed t-tests were performed for comparisons between two groups, and one- or two-way ANOVAs were performed for more than two groups and multiple comparisons. All tests were conducted as two-sided tests with significance levels set as: ns, *p*>0.05; *, *p*<0.05; **, *p*<0.01, ***, *p*<0.001; ****, *p*<0.0001.

## Acknowledgments

We thank Dr. Kazuko Sakata and Dr. Brij B. Singh for constructive discussions, and Dr. Michael P. McDonald for providing 5xFAD breeder mice which had been weaned off the retinal mutant gene. Pictorial illustrations were created with BioRender.com.

## Author contributions

Conceptualization: F-F. L.

Investigation: J. W., L. C., Z. Wang, D. D., G. L., H. Z., S. Z., V. K. B.

Supervision: Z. Wu., W. L., I. B., F-M. Z., J. D.

Writing—original draft: J. W., F-F. L.

Writing—review & editing: J. W., F-F. L.

## Data availability

Raw and processed RNA-seq data used in this study are available at the Gene Expression Omnibus under the accessions GSE253510. Codes used in this study are available from the authors J. W. and Z. Wang. upon request.

