## supplemental figures for "TRPC3 suppression ameliorates synaptic dysfunctions and memory deficits in Alzheimer’s disease"

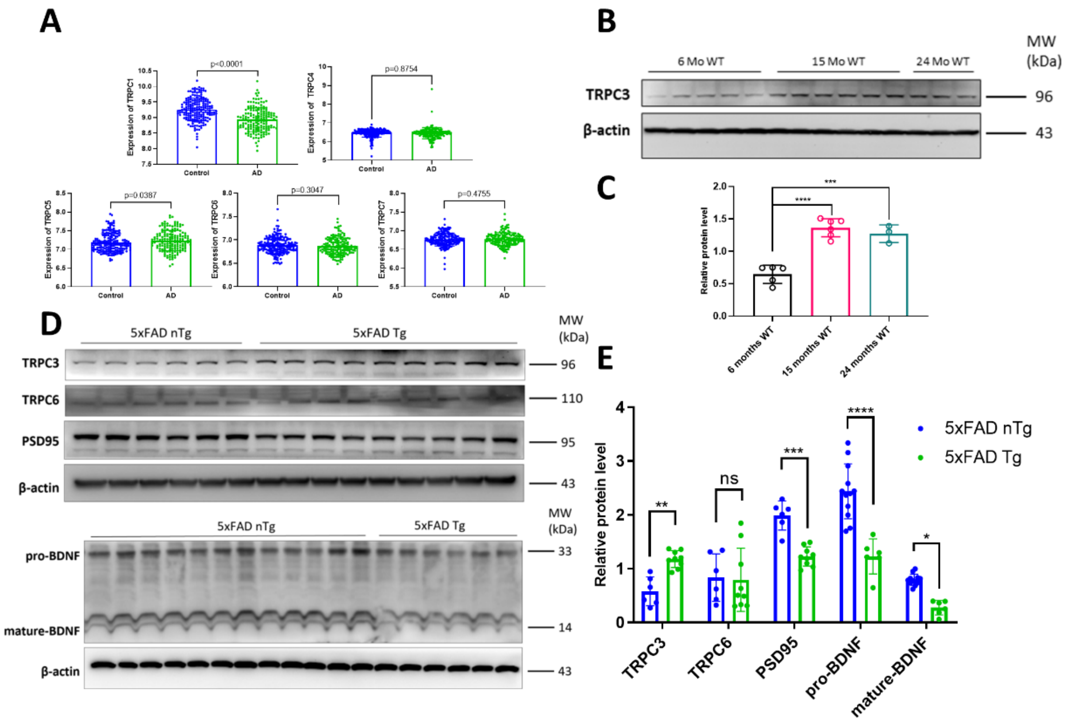


**Fig. S1. A**, The gene expression levels of the remaining TRPC family members based on published microarray database[61]. **B**, TRPC3 protein levels in WT mice at three ages as detected. By Western blots based on three independent experiments. The blots are quantified in **C**. **D**, Hippocampal protein expression levels of TRPC3, TRPC6, and PSD95 in five-month-old 5xFAD mice, with n=6 for nTg and n=9 for Tg. Another cohort of 5xFAD littermates were analyzed for the expression of BDNF, with n=12 for nTg and n=6 for Tg. **E**, Quantification of the blots in panel **D**.

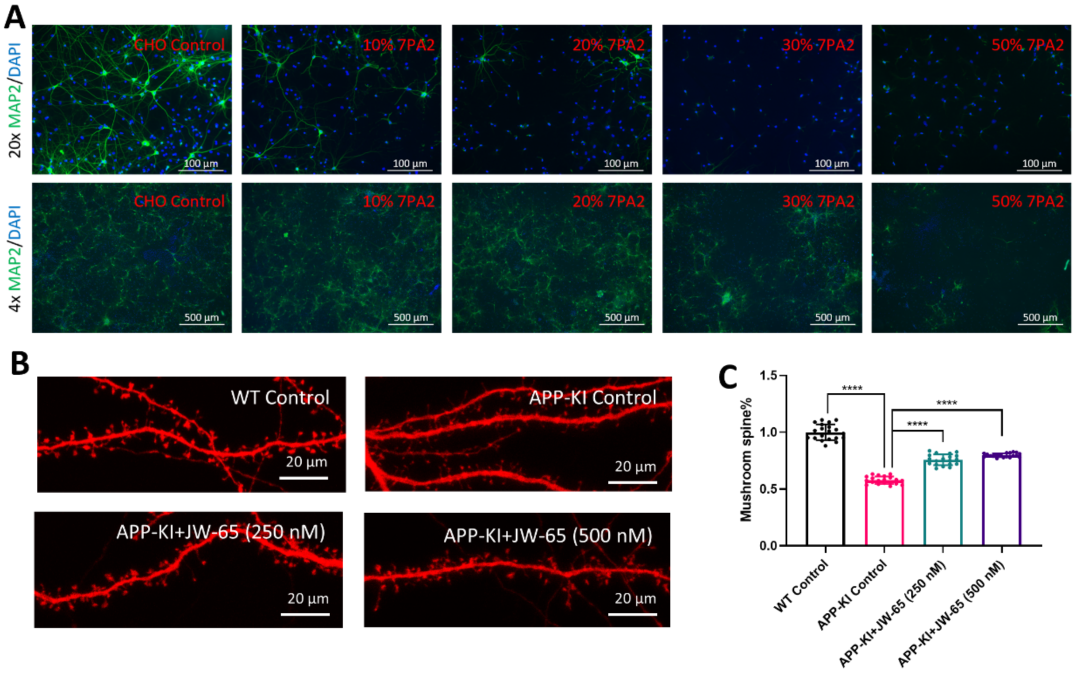


**Fig. S2. Effects of compound treatment on neuronal morphology and gliosis. Immunostaining on rat primary hippocampal culture. A**, Representative microscopic images showing that 7PA2 dose-dependently induces neuronal loss as evidenced by MAP2 staining. **B**, Representative confocal images of the WT and APP-KI hippocampal neurons transfected with TD-Tomato and treated with different concentrations of JW-65 and Pyr3 on DIV15 and fixed at DIV16. **C**, Quantification of the percentage of the mushroom spines based on the images taken in the same experiments of panel **B** (n=18-20 neurons from three batches of cultures).

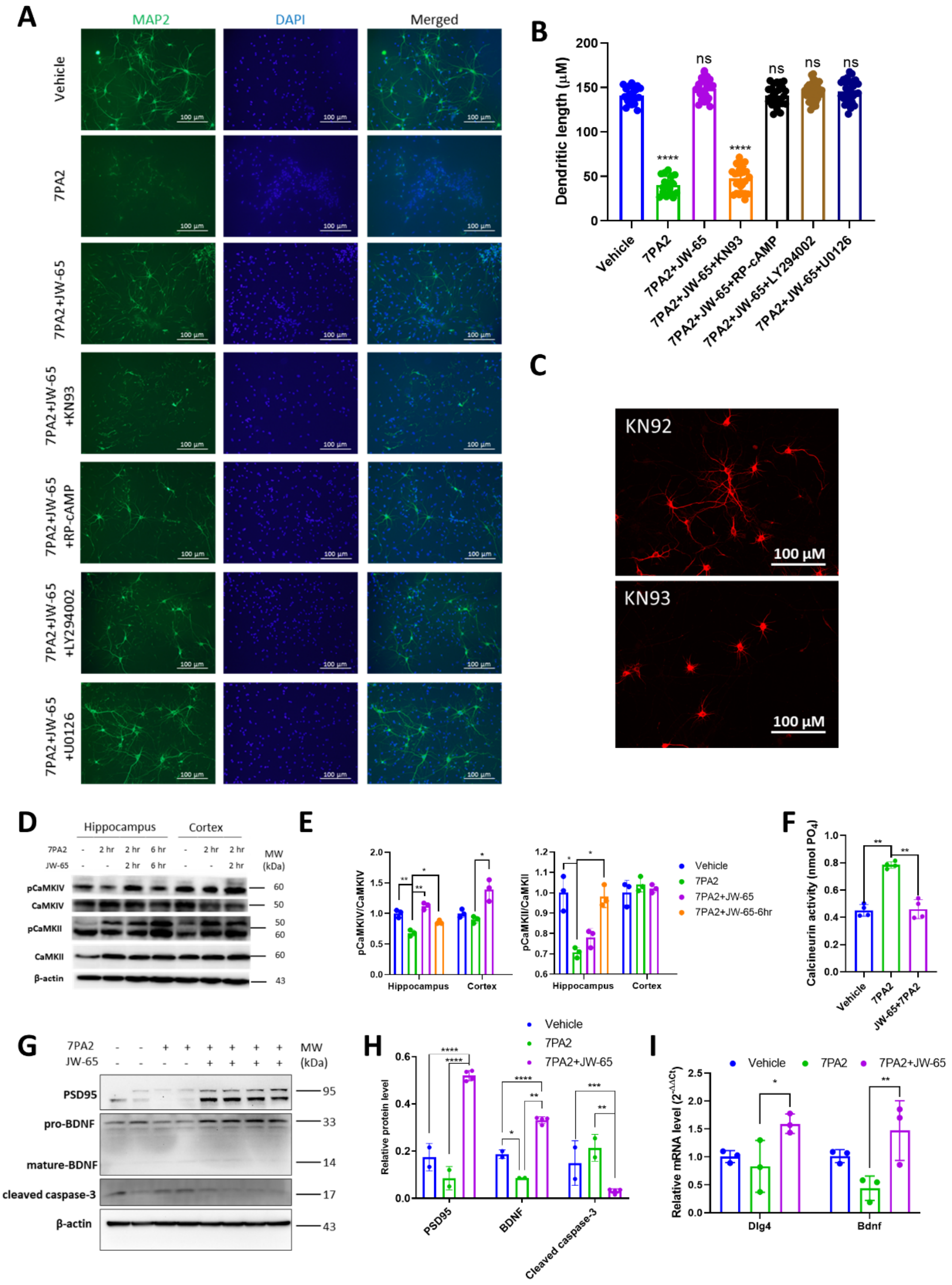


**Fig. S3. Cell signaling pathways underlying neuroprotective JW-65. A**, Representative images show the neuroprotection of JW-65 (500 nM) was counteracted by KN93 (10 μM), a CaMKII inhibitor, but stayed unaffected with the co-treatment of other kinase inhibitors, including RP-cAMP (10 μM), LY294002 (10 μM), and U0216

(10 μM). **B**, Quantification of dendritic length under the co-treatment with different kinase inhibitors. n=18-33. **C**, MAP2 Immunostaining of DIV14 primary neurons treated with KN92 (10 μM) or KN93(10 μM). **D**, Western blots showing the phosphorylation levels of CaMKII and CaMKIV in DVI14 hippocampal neurons and cortex neurons. The blots are quantified in panel **E**. **F**, Quantified graph showing changes in Calcineurin/CaN activity 2 hr after 7PA2 treatment. (n=3) **G**, Western blots showing the expression levels of PSD95, BDNF, and cleaved caspase-3 in DIV14 neurons treated with 7PA2 or in combination with JW-65 (500 nM). The blots are quantified in panel **H**. **I**, mRNA levels of *Dlg4* and *Bdnf* in the same cohort of neuronal samples in panel **G**.

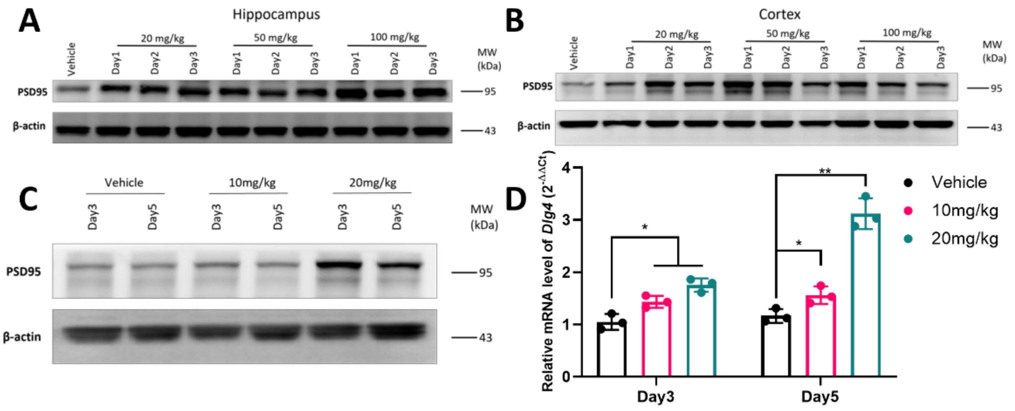


**Fig. S4 Dose response of JW-65 treatment on the expression of synaptic protein PSD95 in WT mice. A-B**, PSD95 expression is enhanced in both the hippocampus and cortex of 4-month-old WT mice post one IP injection of 20/50/100 mg/kg JW-65. **C-D**, Western blot and RT-qPCR indicate IP administration of 20 mg/kg of JW-65 more significantly increased hippocampal PSD95 (*Dlg4*) level compared to 10 mg/kg.

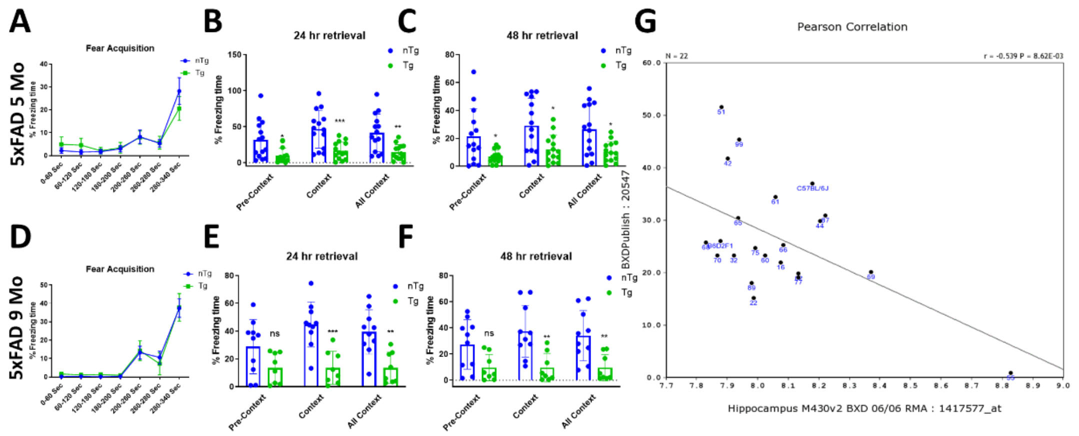


**Fig. S5 TRPC3 expression level is negatively correlated with fear contextual memory. A-C**, Fear contextual training and 24-48 hr fear retrieval test on 5-month-old 5xFAD mice. n=14. **E-F**, Fear contextual training and 24-48 hr fear retrieval test on 9-month-old 5xFAD mice. nTg, n=10. Tg, n=8. **G**, *Trpc3* transcript expression from the hippocampus of 22 recombinant inbred strains (BXD type) revealed a negative relationship between its expression and contextual fear memory (Pearson correlation, r = −0.539, $p = 0.0086$).

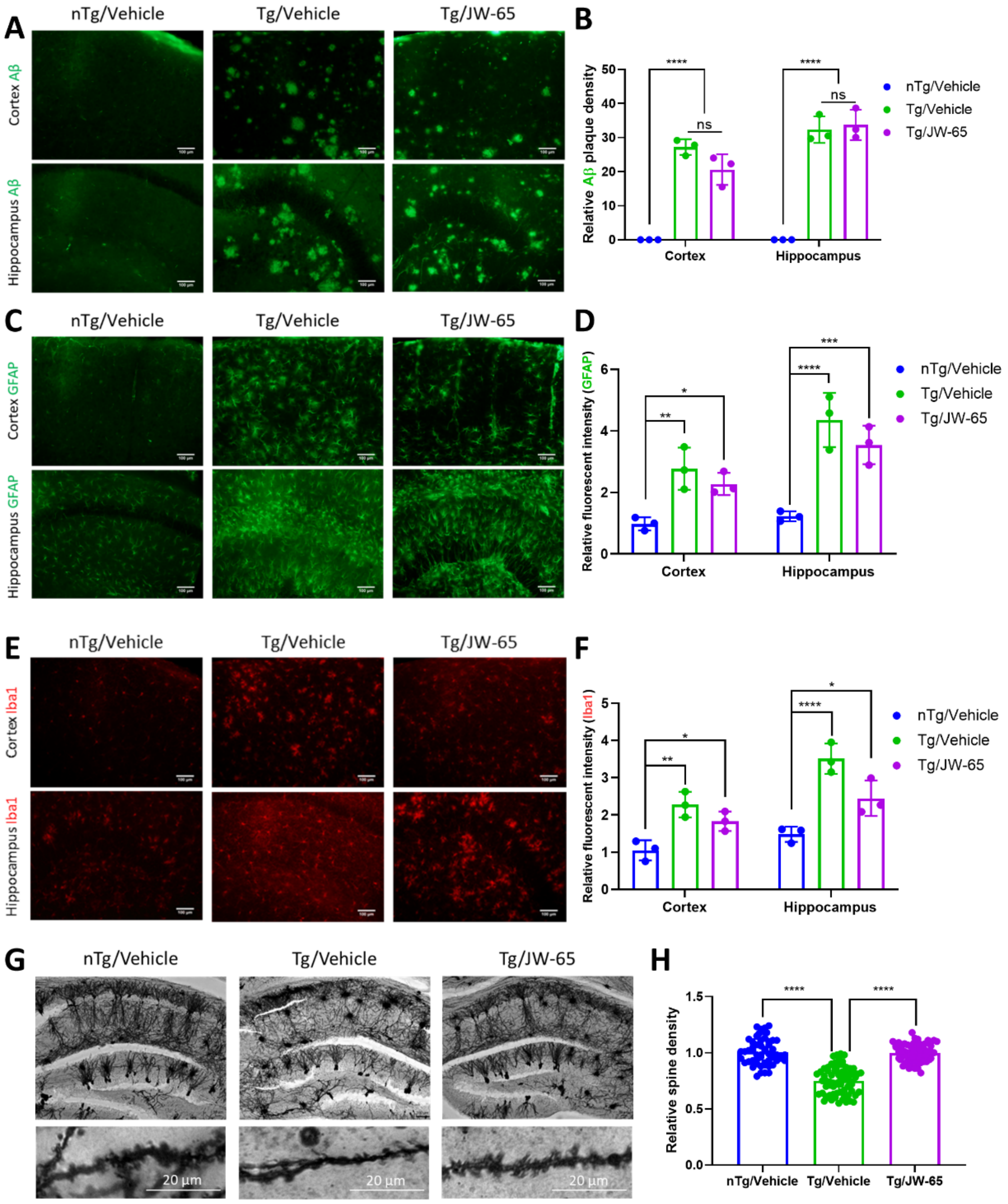


**Fig. S6 JW-65's effects on AD pathological hallmarks in treated 5xFAD mice (n=3 mice each genotype). A-B,** Representative images and quantification of IHC staining of anti-Aβ antibody. **C-D**, Representative images and quantification of IHC staining of microglia in the cortex and hippocampus of 5xFAD mice. Iba1 was used as the marker for microglia. **E-F**, Representative images and quantification of IHC staining of astrocyte in the cortex and hippocampus of 5xFAD mice. GFAP was used as the marker for astrocytes. **G-H**, Representative images and quantification of Golgi staining of the hippocampus in 5xFAD mice.

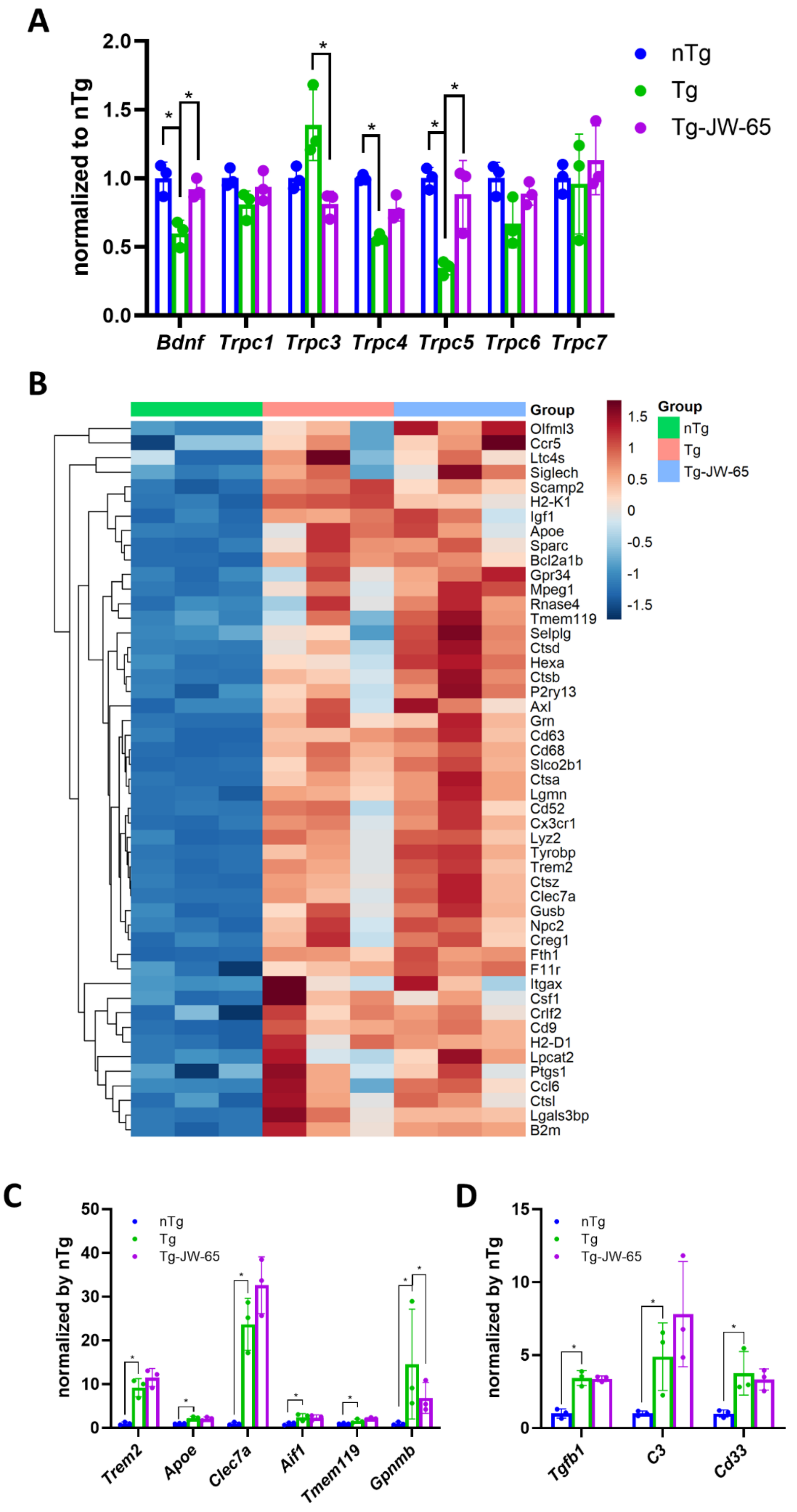
A
normalized to nTg
2.0
1.5
1.0
0.5
0.0
Bdnf
Trpc1
Trpc3
Trpc4
Trpc5
Trpc6
Trpc7
nTg
Tg
Tg-JW-65
B
Group
Olfml3
Ccr5
Ltc4s
Siglech
Scamp2
H2-K1
Igf1
Apoe
Sparc
Bcl2a1b
Gpr34
Mpeg1
Rnase4
Tmem119
Selplg
Ctsd
Hexa
Ctsb
P2ry13
Axl
Grn
Cd63
Cd68
Slco2b1
Ctsa
Lgmn
Cd52
Cx3cr1
Lyz2
Tyrobp
Trem2
Ctsz
Clec7a
Gusb
Npc2
Creg1
Fth1
F11r
Itgax
Csf1
Crlf2
Cd9
H2-D1
Lpcat2
Ptgs1
Ccl6
Ctsl
Lgals3bp
B2m
1.5
1
0.5
0
-0.5
-1
-1.5
Group
nTg
Tg
Tg-JW-65
C
normalized by nTg
50
40
30
20
10
0
nTg
Tg
Tg-JW-65
Trem2
Apoe
Clec7a
Aif1
Tmem119
Gpnmb
D
normalized by nTg
15
10
5
0
nTg
Tg
Tg-JW-65
Tgfb1
C3
Cd33

**Fig. S7. Bulk RNA-seq data from hippocampal tissue. A**, Relative gene expression levels encoding for the TRPC family members and BDNF. **B**, Heatmap of microglia-associated genes. **C-D**, Relative gene expression levels for the microglial DAM and homeostasis genes as well as gene encoding for two complement C3.

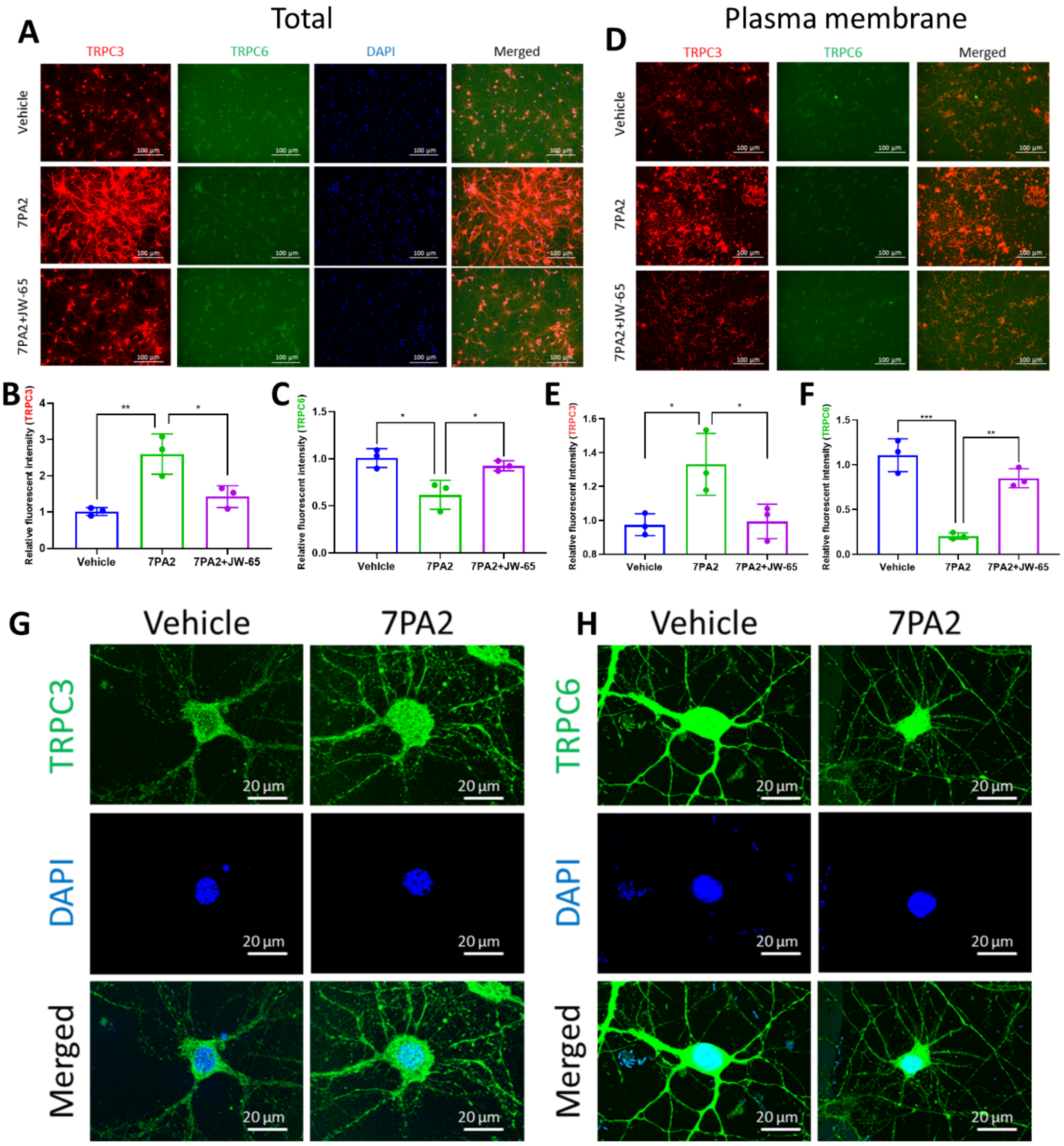


**Fig. S8. AβOs upregulate TRPC3 and downregulate TRPC6 expression in excitatory neurons. A-C**, Representative images of immunocytochemistry of TRPC3 and TRPC6 in neuronal cultures treated with 7PA2 for 4 hr. **D-F**, Representative images of immunocytochemistry performed from cells without permeabilization. **G-H**, Representative confocal images of TRPC3 and TRPC6 showing expressional changes after 7PA2 treatments (4 hr).

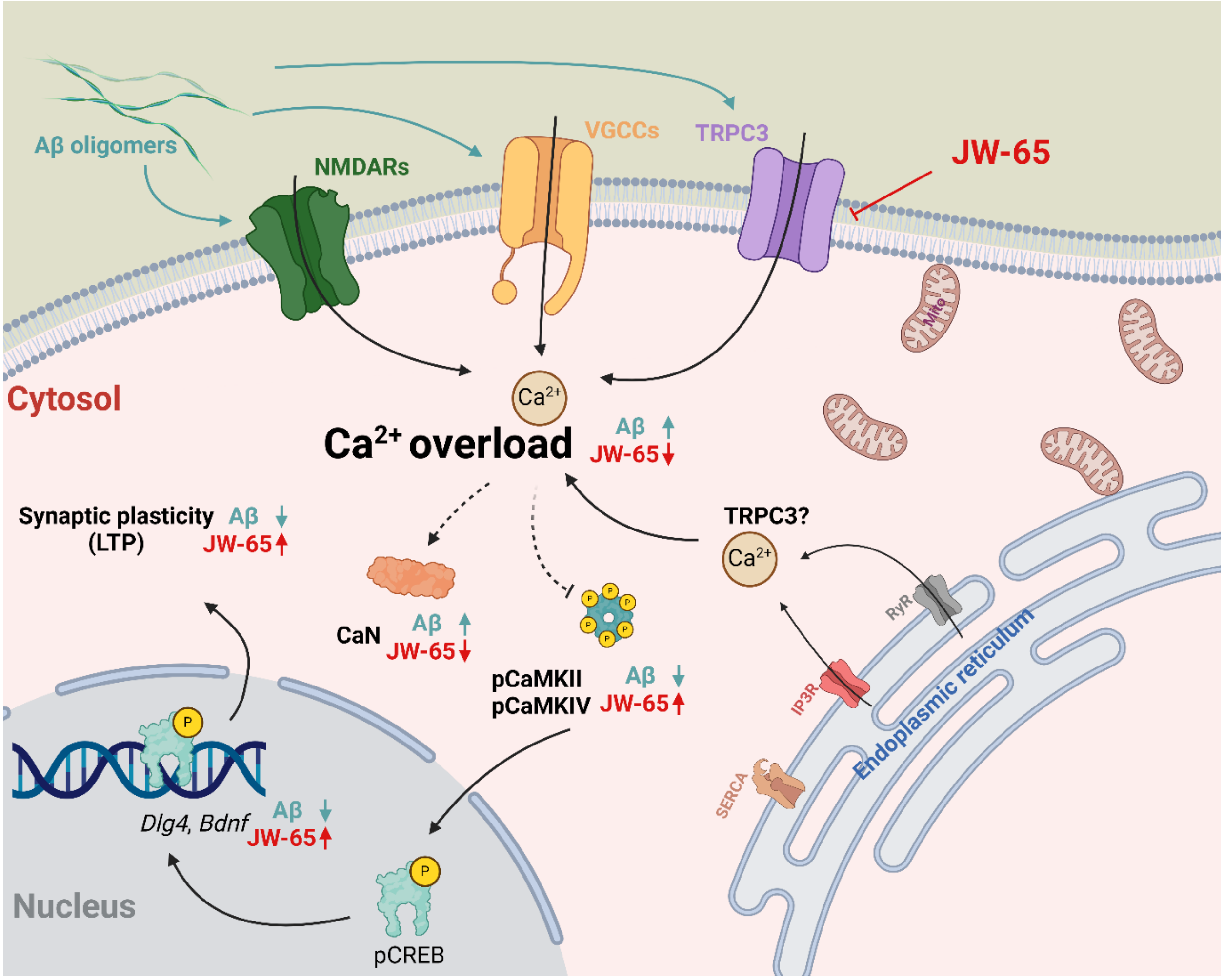


**Fig. S9. Graphical abstract.** Schematic summary of our findings of the novel contribution of TRPC3 in $Ca^{2+}$ signaling in synaptic plasticity and its aberrant roles in mediating $Ca^{2+}$ overload induced by Aβ oligomers. In addition to the well-established ion channels of NMDA receptors (NMDARs) and voltage-gated $Ca^{2+}$ channel (VGCCs), TRPC family member such as TRPC3 also plays significant roles contributing to $Ca^{2+}$ overload through both entry from the plasma membrane and the store-released mechanisms. Of note, the subsequent cell signaling events downstream of the $Ca^{2+}$ overload, including impaired CaMKII/IV activities, along with the downregulated synaptic effector genes (e.g., *Bdnf* and *Dlg4*), as well as the overactivated calcineurin (CaN) can be corrected and restored by TRPC3-selective antagonist compound JW-65. An outstanding question remains as whether TRPC3-mediated $Ca^{2+}$ entry is through SOCE and how TRPC3 is involved in modulating SERCA (ER $Ca^{2+}$-ATPase) expression at the ER membrane and their potential interplay with the inositol 1,4,5-triphosphate receptors ($InsP_3Rs$) or ryanodine receptors (RyRs) in $Ca^{2+}$ release from ER also warrants further investigation.

**Table S1. List of primers used in this research.**

| Primer | sequence |
|---|---|
| *mTrpc3*, forward | TTAATTATGGTCTGGTTCTTGG |
| *mTrpc3*, reverse | TCCACAACTGCACGATGTACT |
| *mTrpc6*, forward | GCAGCTGTTCAGGATGAAAC |
| *mTrpc6*, reverse | TTCAGCCCATATCATGCCTA |
| *mDlg4*, forward | TCTGTGCGAGAGGTAGCAGA |
| *mDlg4*, reverse | CGGATGAAGATGGCGATAG |
| *mBdnf*, forward | AATTAAGCTTCCAATCGAAGCTCAACCG |
| *mBdnf*, reverse | AATTGAAATTCTCCACACAAAGCTCTCGGA |
| *mGapdh*, forward | GCAAATTCAACGGCACAG |
| *mGapdh*, reverse | CTCGCTCCTGGAAGATGG |
| *rTrpc3*, forward | CCACATGCAGTGAGACTTTGACTC |
| *rTrpc3*, reverse | AGGCCAACCTTGGGATCATT |
| *rTrpc6*, forward | AGAAATTTGGAATTTTGGGAAGTC |
| *rTrpc6*, reverse | TCCTTATCAATCTGGGCCTGC |
| *rDlg4*, forward | GGCACACAAGTTCATTGAGG |

| Primer | sequence |
|---|---|
| *rDlg4*, reverse | GAGACATCGAGGATGCAGTG |
| *rBdnf*, forward | AGCGCGAATGTGTTAGTGGT |
| *rBdnf*, reverse | GCAATTGTTTGCCTCTTTTTCT |
| *rbActin*, forward | CCCGCGAGTACAACCTTCT |
| *rbActin*, reverse | CGTCATCCATGGCGAACT |

**Table S2. List of antibodies used in this research.**

| Antibody | Cat# |
|---|---|
| TRPC3 | ACC-016 |
| TRPC6 (WB) | ACC-017 |
| TRPC6 (ICC) | ab105845 |
| PSD95 | 51-6900 |
| BDNF | ANT-010 |
| pCaMKII | 12716 |
| CaMKII | 3362 |
| pCaMKIV | sc-28443-R |
| CaMKIV | sc-166156 |
| Cleaved caspase-3 | 9611 |

| **Antibody** | **Cat#** |
|---|---|
| β-actin | A2228 |
| Synapsin 1 | A-6442 |
| Iba1 | 019-19741 |
| GFAP | G3893 |
| Goat anti-Mouse HRP | 31430 |
| Goat anti-Rabbit HRP | 31460 |
| Goat anti-Mouse Alexa Fluor Plus 488 | A32723 |
| Goat anti-Rabbit Alexa Fluor Plus 594 | A21207 |
